# Mapping Mesolimbic Dopamine Projections to Subcortical Circuits Underlying Adaptive Behavior: A Human 7T Diffusion Tractography Atlas

**DOI:** 10.64898/2026.09.22.753583

**Authors:** Blake L. Elliott, Ranesh Mopuru, Linda J. Hoffman, Josiah K. Leong, Daniel Zweben, Nora D. Volkow, Ingrid R. Olson, Vishnu P. Murty

**Author notes:** Correspondence: Blake L. Elliott. Blake L. Elliott and Ranesh Mopuru contributed equally to this work.

## Abstract

The mesolimbic dopamine system supports broad aspects of adaptive behavior including salience, learning, memory, and decision making, yet the structural organization supporting these functions remains incompletely mapped in humans. Using high-resolution 7T diffusion magnetic resonance imaging (up to N=173), we reconstructed seven bilateral ventral tegmental area (VTA)–subcortical pathways spanning direct mesolimbic projections and the hippocampus–VTA (HPC–VTA) loop regulatory circuit. Projection-endpoint analyses revealed long-axis and subfield organization of VTA–hippocampal pathways and superior-inferior VTA–nucleus accumbens (NAc) topography. Consistent with the HPC–VTA loop framework, neurite density covaried across mesolimbic pathways but not the arcuate fasciculus, demonstrating circuit-level organization. Nodewise analyses identified localized associations between pathway microstructure and delay discounting across VTA–hippocampal, VTA–NAc, and VTA– amygdala pathways. Together, these findings parse the human mesolimbic system into distinct pathways supporting adaptive behavior. The probabilistic atlas, MesoConnect, is freely available with tutorials to facilitate reconstruction and investigation of circuits in independent datasets.

## Introduction

Adaptive behavior requires more than detecting rewards. An organism must identify events that matter, learn their consequences, bind them to context for selection and to motivate action, and update future behavior. Mesolimbic circuits contribute to diverse behaviors including motivational salience detection, appetitive and aversive learning, behavioral activation, motivated memory, novelty processing, and adaptive choice. ^1–5^ The mesolimbic dopamine system originates from the ventral tegmental area (VTA) and ascends through the medial forebrain bundle (MFB) to support these functions.^6–8^ Disruption of this circuitry is implicated in a wide range of psychiatric and neurological disorders.^9–11^ Although these pathways are well characterized in animal models, their pathway-resolved organization remains poorly mapped in humans.

The VTA is anatomically and functionally heterogeneous. Classical axonal tracing studies show overlapping but systematic projection topography across medial-lateral, dorsal-ventral, and anterior-posterior axes.^12–14^ Projection target also relates to the molecular, electrophysiological, and input-output properties of VTA dopamine neurons.^15–18^ Defining VTA neurons by projection target reveals distinct populations carrying signals related to reward, salience, sensation, and action.^19–21^ Human diffusion tractography and cadaveric microdissection support similar connectivity-dependent organization, although human studies remain limited and have often examined connectivity gradients across the larger VTA/substantia nigra (SN) rather than pathways arising specifically from the VTA.^22–25^ These findings suggest a graded topology that biases information flow without imposing a fixed one-pathway, one-function mapping. Accordingly, we organized the present study around three related functions: motivated learning and salience, represented by VTA–NAc and VTA–amygdala pathways; motivated memory, represented by VTA–anterior and –posterior HPC pathways; and regulation of motivated behavior, represented by the polysynaptic HPC–NAc–VP and VP–VTA circuit. The VTA–HPC and HPC–NAc–VP–VTA pathways form complementary components of the HPC–VTA loop: an upward VTA–HPC arc that modulates hippocampal plasticity and a downward HPC–NAc–VP– VTA arc that regulates VTA population activity.^3,26^

Within the motivated learning and salience system, VTA connections with the NAc and amygdala provide complementary routes through which motivational significance can shape learning and action. Phasic dopamine responses can signal reward prediction errors, while NAc dopamine also contributes to incentive salience, or motivational “wanting,” and behavioral activation, or the invigoration of goal-directed action.^1,5,27,28^ Animal tracing shows medial-lateral organization, with medial VTA dopamine neurons preferentially projecting to medial NAc territories and lateral VTA neurons projecting to lateral NAc territories,^13,15,16,29^ while functional signals vary by projection target and behavioral context.^17,30–32^ Human studies link VTA–NAc pathway structure to impulsivity and substance use and identify distinct inferior and superior trajectories on opposite sides of the anterior commissure.^23,24^ VTA–amygdala circuitry provides a complementary pathway for affective salience and associative learning, with converging evidence that midbrain dopamine input and amygdala function contribute to appetitive, aversive, and reinforcement learning.^33–35^

Motivated memory centers on interactions between the VTA and HPC.^4^ Animal studies identify direct dopaminergic projections from the VTA to the HPC, with denser innervation of ventral hippocampal territories, particularly the subiculum and adjacent CA1, while also demonstrating projections to dorsal hippocampus.^36–39^ Hippocampal dopamine promotes long-term potentiation and memory persistence, and VTA–HPC dopamine manipulations modulate hippocampal plasticity and contextual memory.^37,40–42^ Human structural evidence remains sparse, but studies link midbrain–HPC tract density to individual differences in monetary gain- and loss-motivated memory encoding, demonstrate an anterior bias in VTA–HPC structural connectivity, and identify VTA connections with the dorsal dentate gyrus using microdissection-guided tractography.^22,25,43^ These findings motivated separate reconstructions of the VTA–Anterior HPC and VTA–Posterior HPC pathways.

Beyond these VTA–centered pathways, a polysynaptic HPC–NAc–VP–VTA circuit regulates motivated behavior by controlling VTA population activity. Excitatory HPC output to the NAc suppresses VP inhibition of the VTA, recruiting otherwise silent VTA dopamine neurons into spontaneous firing. By controlling the size of this spontaneously active population, HPC input regulates how many dopamine neurons are available for subsequent phasic responses, effectively setting the gain of the VTA dopamine system.^26,44^ Studies implicate components of this circuit in depression-like behavior, substance-related learning and seeking, and HPC-driven dopamine dysregulation in models of psychosis.^26,45–47^ Human diffusion imaging likewise links stronger HPC–ventral striatal structural connectivity to greater binge-drinking frequency.^48^ Because this downward arc is polysynaptic, we reconstructed HPC–NAc–VP, with the NAc serving as an intermediate waypoint between HPC and VP, and reconstructed VP–VTA separately.

Here, we used high-resolution 7T diffusion magnetic resonance imaging from the Human Connectome Project (HCP) to create MesoConnect, a probabilistic atlas of seven bilateral pathways: VTA–Inferior NAc, VTA–Superior NAc, VTA–Amygdala, VTA–Anterior HPC, VTA–Posterior HPC, HPC–NAc–VP, and VP–VTA (Figures 1-2). We pursued four goals. First, we reconstructed anatomically constrained pathways in participant-native space and generated population probability maps. Second, we quantified pathway geometry and endpoint topology within the HPC, NAc, and VTA. Third, we tested whether neurite density index (NDI) covaried across the circuit’s constituent pathways relative to a spatially proximal non-mesolimbic comparison tract, as evidence of coordinated circuit organization. Finally, as a proof of concept for relating pathway-resolved microstructure to individual differences in motivated behavior, we tested NDI associations with delay discounting, a task that broadly engages mesolimbic circuitry by integrating valuation, future-outcome representation, salience, memory, and action selection.^49–51^ We predicted that the atlas would reveal reproducible connectional heterogeneity and provide an open structural framework for testing how distinct mesolimbic pathways operate separately and together in adaptive and maladaptive behavior. The MesoConnect atlas files and analysis code are available through GitHub (https://github.com/blelliott23/MesoConnect), and step-by-step documentation and tutorials are available on the MesoConnect website (https://mesoconnect.vercel.app).

**Figure 1.**
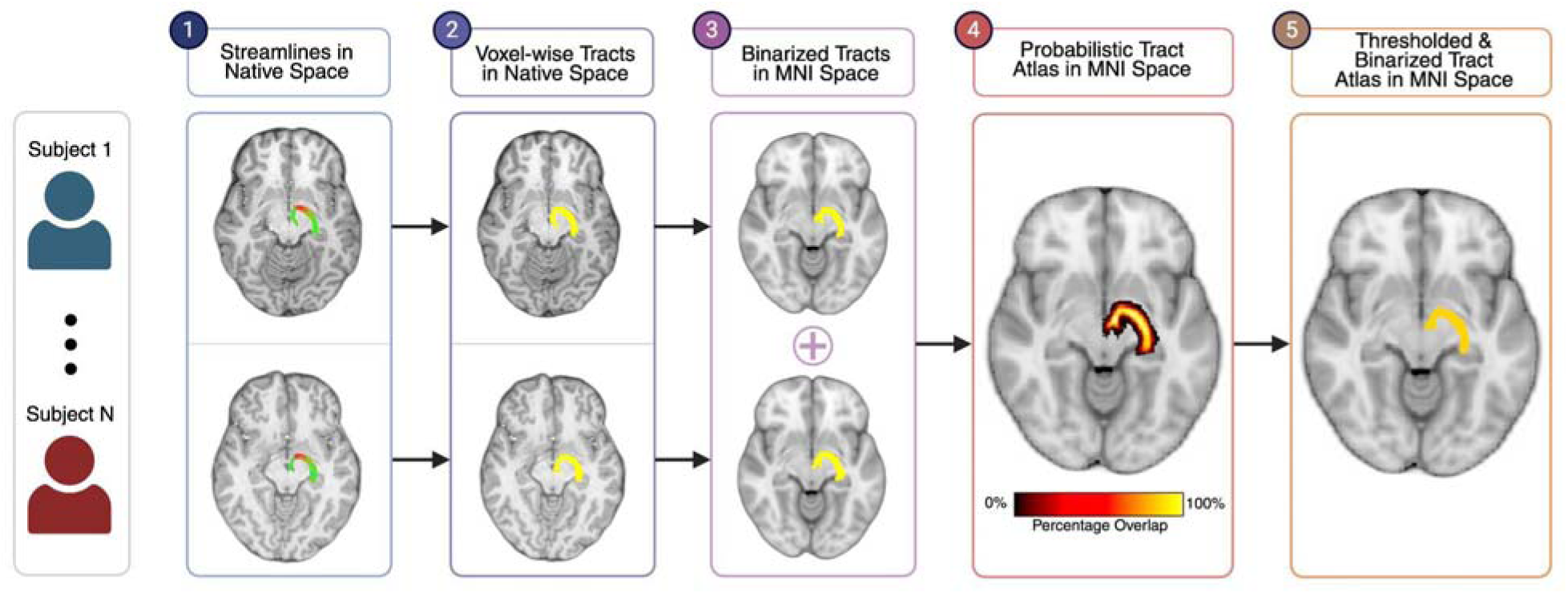
MesoConnect Atlas-generation workflow. Subject-native tractograms were converted to binary maps, transformed to Montreal Neurological Institute (MNI) standard space, aggregated across participants, and thresholded at multiple participant-overlap levels to produce group probabilistic atlases.

**Figure 2.**
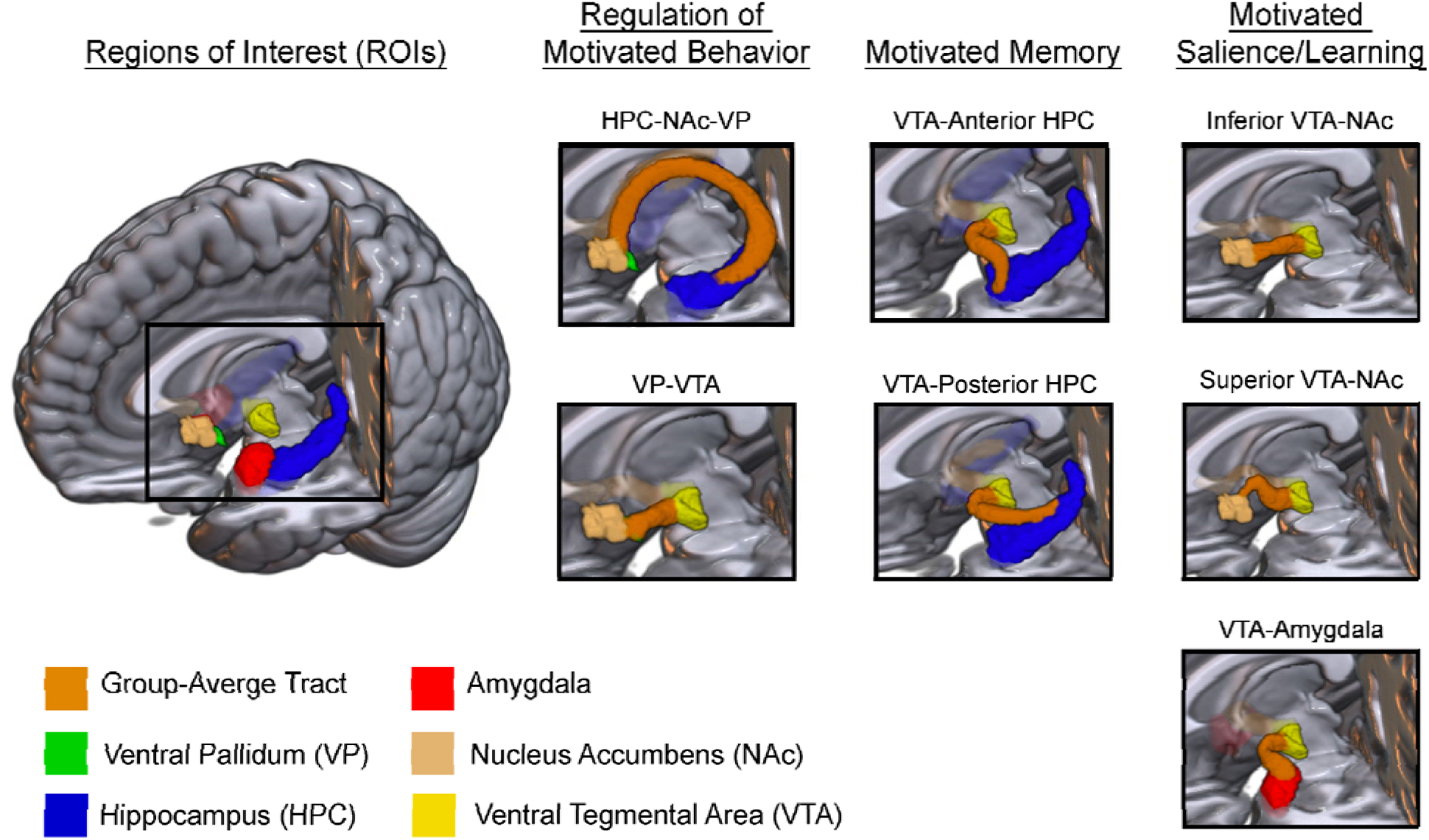
MesoConnect Mesolimbic Connectivity Atlas. Group-average pathways and anatomical regions of interest are organized into three proposed functional groupings: regulation of motivated behavior (HPC–NAc–VP and VP–VTA), motivated memory (VTA–anterior HPC and VTA–posterior HPC), and motivated salience/learning (inferior VTA–NAc, superior VTA–NAc, and VTA–amygdala). The motivated-memory tracts form the VTA–centered “upward arc,” whereas the regulation-of-motivated-behavior pathways represent separable components of the “downward arc” of the HPC–VTA loop circuit. HPC–NAc–VP was reconstructed with the NAc as an intermediate waypoint between the HPC and VP; VP–VTA was reconstructed separately. Orange indicates the group-average tract (50% probability threshold); region colors are defined in the figure.

## Results

### Atlas reconstruction, coverage, and cross-participant agreement

All seven bilateral pathways were reconstructed in most of the 173 tractography-eligible participants (Figure 2; Table 1), with successful bilateral reconstruction in 166–173 participants per pathway (96.0%–100.0%). The VTA-centered atlas resolved pathways to anterior and posterior HPC territories, the amygdala, and superior and inferior NAc territories, whereas the “downward-arc” atlas resolved the HPC–NAc–VP and VP–VTA components of the regulatory circuit (Figures 2–3). Participant mean streamline length was stable within pathways, with low between-participant variability across all pathway and hemisphere combinations (Supplementary Table S8).

**Figure 3.**
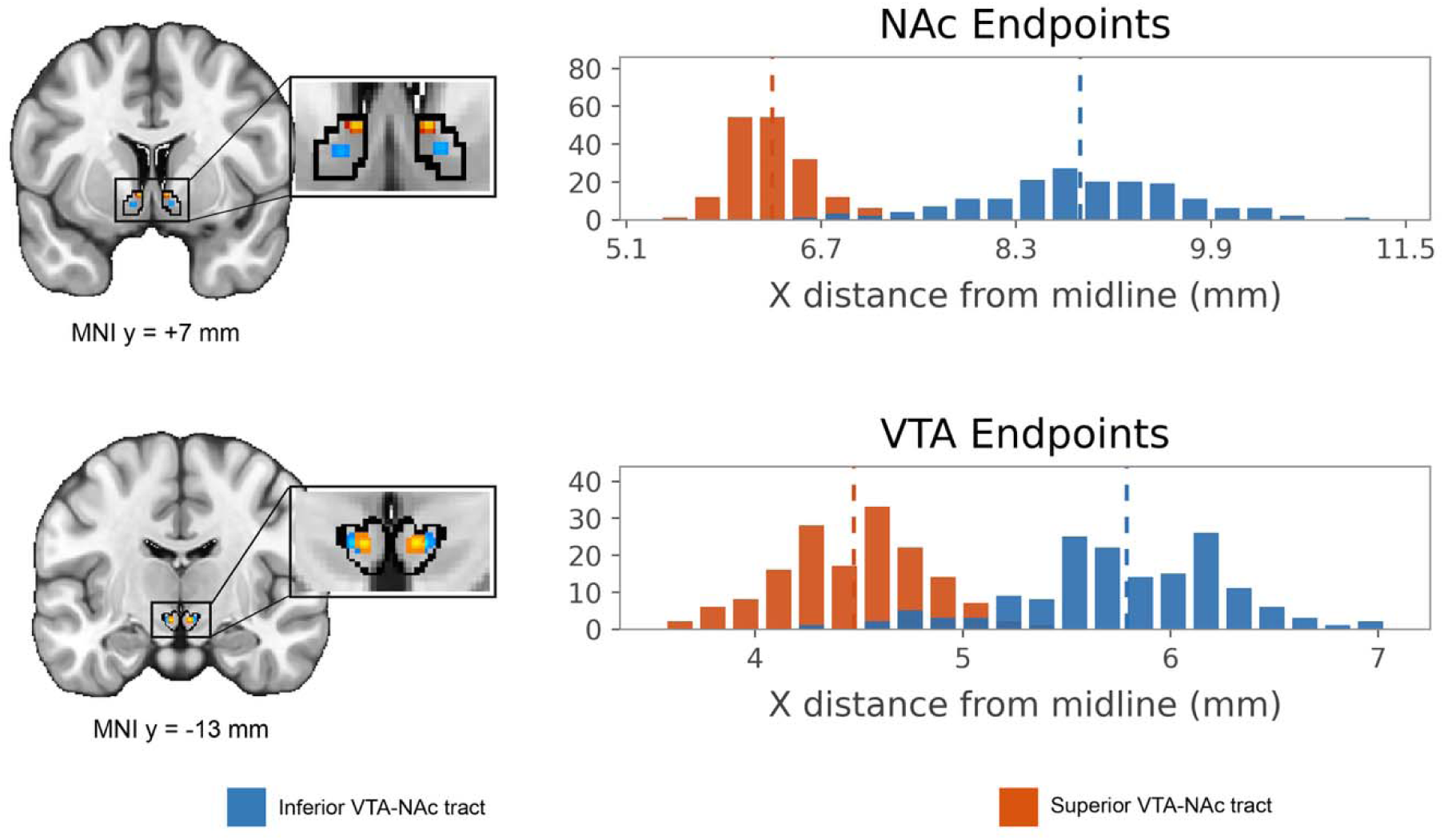
Endpoint topography of the inferior and superior VTA–NAc tracts. Endpoint-density maps and histograms of distance from the midline show separation of the two pathways at both the NAc and VTA ends of the reconstruction. At the NAc, superior-pathway endpoints were more medial, anterior, and dorsal than inferior-pathway endpoints. Coordinates were collapsed bilaterally by averaging |x|, y, and z.

**Table 1.**
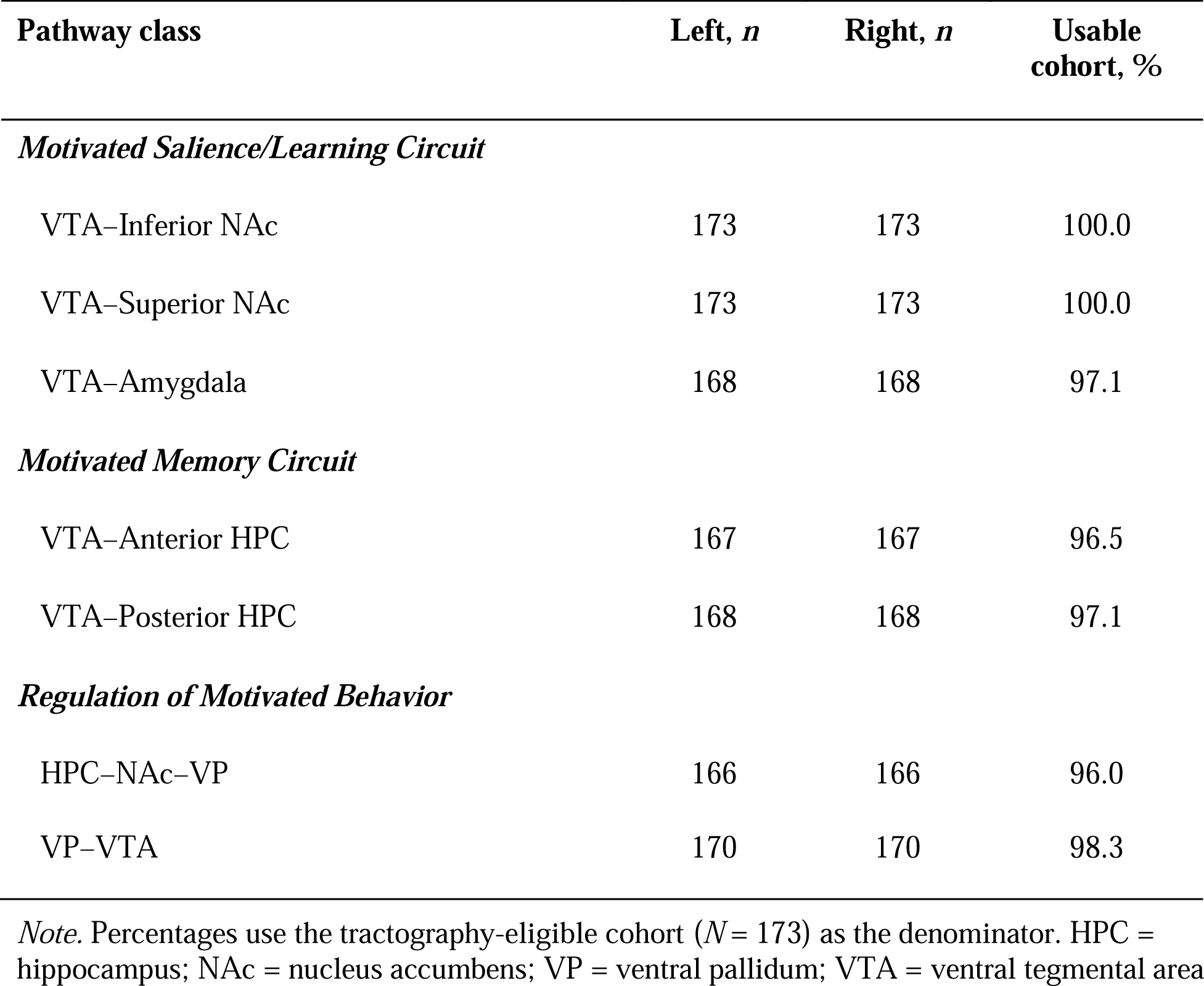
Pathway Classes and Valid Group-Atlas Maps.

Fixed-atlas agreement was highest at the 25% and 50% overlap thresholds (Supplementary Table S3). At 50%, pooled median ordinary Dice was .687, dilated Dice .761, 95th-percentile Hausdorff distance was 1.819 mm, bundle overlap was .632, and overreach was .162. As expected, the 25% atlas increased participant coverage with greater overreach, whereas the 75% atlas reduced overreach at the cost of coverage. We therefore use the 50% map as the conservative primary atlas, with 25% and 75% maps providing more sensitive and stricter consensus representations, respectively.

### Motivated learning and salience circuit: VTA–NAc and VTA–Amygdala pathways

The inferior and superior VTA–NAc tracts were separable at both endpoints (Figure 3). At the NAc, the inferior pathway approached a more lateral, posterior, and inferior territory than the superior pathway (superior-minus-inferior: −2.53 mm from the midline, +1.00 mm anterior-posterior, +4.43 mm superior-inferior; Hotelling’s T² = 10,986.55, F(3, 169) = 3,619.35, p < .001). At the VTA, the superior pathway approached a more medial, slightly anterior, and slightly inferior territory (−1.31, +0.39, and −0.13 mm, respectively; Hotelling’s T² = 1,628.92, F(3, 153) = 535.97, Holm-adjusted p = 4.49 × 10 ; Supplementary Table S5B). This medial- to-medial and lateral-to-lateral organization parallels animal VTA–NAc projection topography and demonstrates reproducible organization at both ends of the reconstruction.

VTA–amygdala maps were available bilaterally in 168 participants. Qualitative slice-wise inspection showed spatial correspondence between the group VTA–amygdala pathway and the published VTA–temporal reconstruction of Skandalakis et al.^25^ The reconstructed pathway followed a similar course from the VTA toward the amygdala, and its anatomical localization was consistent with landmarks in the MNI-referenced, histology-based ex vivo human brain atlas of Mai et al.^52^ Together, correspondence with cadaveric microdissection and histology-based atlas landmarks supports the anatomical plausibility of the reconstructed VTA–amygdala pathway.

### Motivated memory circuit: Anterior and posterior VTA–HPC pathways

Exact subfield overlap was lower for VTA–anterior HPC than VTA–posterior HPC endpoints because the Harvard-Oxford HPC reconstruction mask was slightly larger than participant-specific segmentations. A one-voxel dilation recovered contact for 178 of 179 initially nonoverlapping maps, yielding near-complete final contact across pathways (Figure 4; Supplementary Table S6A). In the final-contact sample, VTA–anterior HPC endpoints were almost exclusively CA1 dominant and localized to the hippocampal head, whereas VTA–posterior HPC endpoints were more frequently dentate-gyrus dominant and localized to the hippocampal body. CA1 was dominant in 99.4% of left and 98.8% of right anterior maps; dentate gyrus was dominant in 72.6% of left and 64.9% of right posterior maps, approximately 96% of whose contact voxels were located in the hippocampal body. Route differences were pronounced for head-versus-body dominance, head-contact proportion, and dominant-subfield distribution (all pooled p ≤ 1.42 × 10□¹³□; Figure 4; Supplementary Table S6, Panels B–D). These findings demonstrate distinct tractography-defined endpoint territories but do not establish histological termination within particular hippocampal subfields.

**Figure 4.**
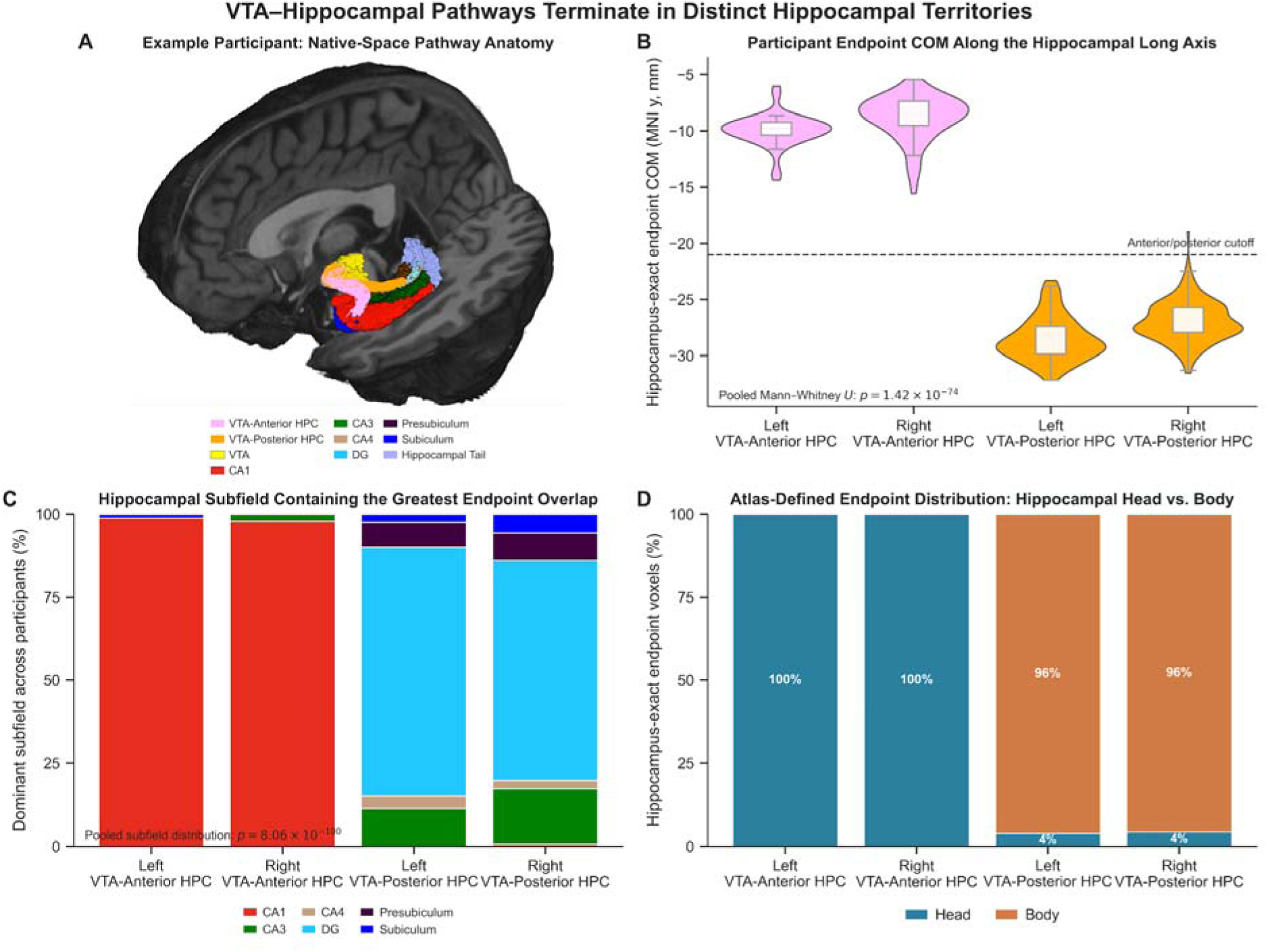
VTA–Anterior HPC and VTA–Posterior HPC approach distinct hippocampal territories. (A) Example native-space pathways and hippocampal subfields. (B) Endpoint center of mass along the hippocampal long axis. (C) Dominant overlapping subfield. (D) Head-versus-body distribution of exact-overlap endpoint voxels. Endpoints index tractography approaches to the region of interest, not histological terminations.

Qualitative slice-wise inspection and three-dimensional rendering showed strong correspondence of both the VTA–Anterior HPC and VTA–Posterior HPC pathways with the cadaveric VTA–temporal microdissection reconstruction of Skandalakis et al.^25^ The VTA–Anterior HPC pathway corresponded to the component grouped with fibers reaching the amygdala and continuing into the entorhinal cortex, whereas the VTA–Posterior HPC pathway corresponded to the component extending toward the posterior hippocampus and dorsal dentate gyrus. Concordance with the cadaveric reconstruction and cytoarchitectonic landmarks in the MNI-referenced, histology-based ex vivo human brain atlas of Mai et al.^52^ further supported the anatomical plausibility of both VTA–HPC routes and their differentiated territories along the hippocampal long axis.

### VTA topography across pathways

The five direct VTA–centered pathways differed systematically in where they exited the VTA (Figure 5; Supplementary Table S5). In 156 participants with complete bilateral observations, endpoint centers of mass differed along all three axes: distance from the midline, F(2.78, 430.37) = 517.55, p < .001, generalized η² = .489; anterior–posterior position, F(2.84, 439.56) = 298.82, p < .001, generalized η² = .561; and superior–inferior position, F(3.13, 485.38) = 93.32, p < .001, generalized η² = .226. All 10 paired three-coordinate pathway contrasts survived Holm correction (Supplementary Table S5, Panel B). To quantify how much pathway identity could be recovered from VTA endpoint location alone, leave-one-subject-out linear discriminant analysis classified the five pathway classes with 71.92% accuracy (95% participant-level bootstrap confidence interval [69.23%, 74.62%]; chance = 20%; Figure 5; Supplementary Table S5). Thus, VTA endpoint location contained substantial information about pathway identity.

**Figure 5.**
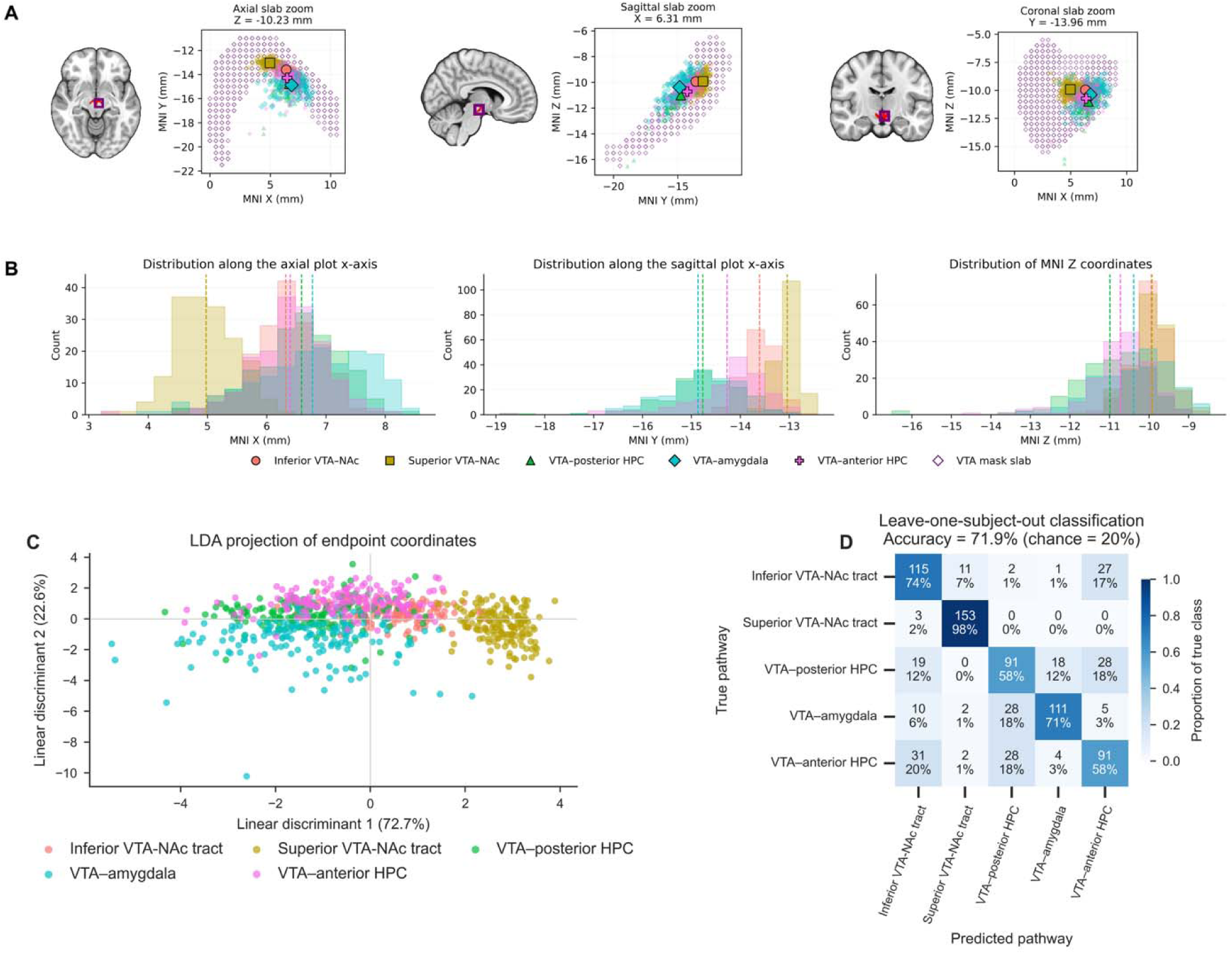
VTA Endpoint Topography and Pathway Classification Across the Five Direct Pathway Classes. (A) Slab views show the spatial organization of pathway-specific VTA-adjacent endpoints in axial, sagittal, and coronal planes. (B) Corresponding coordinate distributions show endpoint variation along the three spatial axes. (C) Projection of distance from the midline x, y, and z endpoint coordinates onto the first two full-sample linear discriminants. (D) Leave-one-subject-out confusion matrix across 156 participants and five pathway classes. Overall classification accuracy was 71.92% (561/780 observations; chance = 20%).

### Regulation of motivated behavior: “downward arc”

The two downward-arc components, HPC–NAc–VP and VP–VTA, were reconstructed separately rather than forcing a continuous HPC–to–VTA streamline across the polysynaptic circuit. Consistent with known anatomy, HPC–NAc–VP followed the fornix and descended through its precommissural columns to the NAc before continuing to the VP.

### Circuit-level covariation with microstructure

To test whether microstructural variation was coordinated across mesolimbic pathways, we calculated partial correlations between bilateral mean whole-tract NDI for all 21 pathway pairs (Figure 6A; Supplementary Table S4). We additionally tested correlations between each mesolimbic pathway and the arcuate fasciculus as a non-mesolimbic comparison and repeated the analyses controlling for whole-white matter (WM) NDI to assess whether associations reflected global white-matter variation. NDI covaried broadly across the mesolimbic pathway set, with 20 of 21 pairwise associations surviving FDR correction. The strongest relationships linked VTA–Anterior HPC with VTA–Amygdala (r = .783), VTA–Posterior HPC with VTA–Amygdala (r = .707), and VTA–Anterior HPC with VTA–Posterior HPC (r = .696). Consistent with the proposed regulatory architecture of the downward arc, VP–VTA covaried with all six other mesolimbic pathways, while HPC–NAc–VP covaried with four of the five VTA–centered pathways; only its association with VTA–Inferior NAc did not survive correction (r = .154, q = .0534). In contrast, none of the seven arcuate-mesolimbic correlations survived correction.

**Figure 6.**
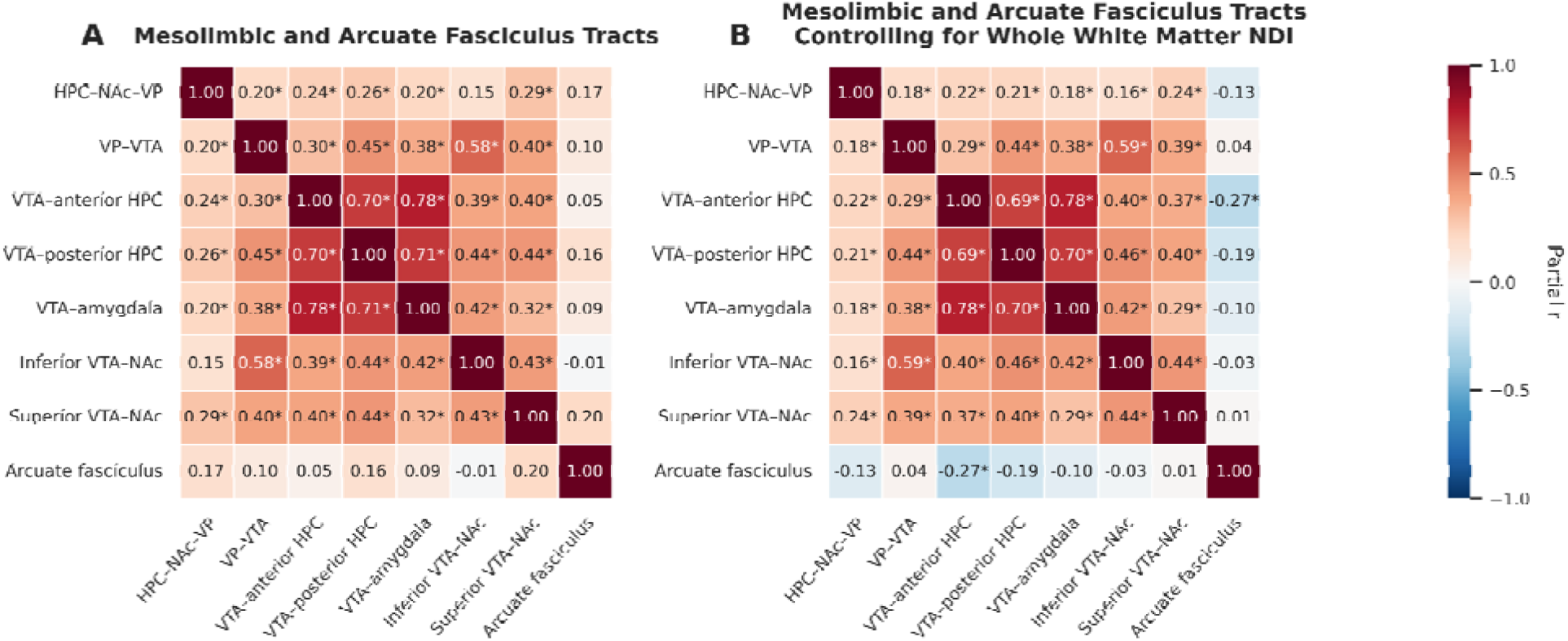
Circuit-level covariation in bilateral mean whole-tract neurite density index. (A) Partial Pearson correlations after adjustment for sex, age, intracranial volume, and handedness. (B) Partial Pearson correlations after additional adjustment for whole-white-matter neurite density index. The arcuate fasciculus is the nonmesolimbic comparison. Benjamini-Hochberg false-discovery-rate correction was applied; asterisks indicate q < .05. Exact statistics are reported in Supplementary Table S4.

Controlling for whole-WM NDI left the positive mesolimbic pattern largely unchanged (Figure 6B; Supplementary Table S4), with all 21 mesolimbic pathway pairs surviving FDR correction. Notably, the HPC–NAc–VP association with the inferior VTA–NAc tract became significant after whole-WM adjustment (r = .158, n = 164, q = .0470). Arcuate-mesolimbic associations were generally negative after adjustment, with a significant negative association emerging with VTA–Anterior HPC. Thus, the persistence of positive covariance within the mesolimbic network, alongside the absence or reversal of corresponding associations with the comparison tract, argues against the pattern being explained by global microstructural variation alone.

### Individual differences in adaptive choice: Delay discounting

To determine whether individual differences in behavior were associated with spatially localized variation in mesolimbic microstructure, we tested nodewise relationships between delay discounting and NDI along each pathway using cluster-mass correction. All five VTA– centered pathways were evaluated bilaterally. Nodewise analyses identified four clusters associated with steeper discounting (Figure 7; Supplementary Table S7): left inferior VTA–NAc (nodes 0-47, family-wise error rate corrected p = .0106), right superior VTA–NAc (nodes 19-46, p = .0191), left VTA–amygdala (nodes 8-29, p = .0392), and left VTA–posterior HPC (nodes 0-17, p = .0434). Because discounting scores were sign-reversed, positive coefficients indicate higher NDI with steeper discounting. These results localize behavioral associations across VTA– NAc, VTA–amygdala, and VTA–HPC pathways.

**Figure 7.**
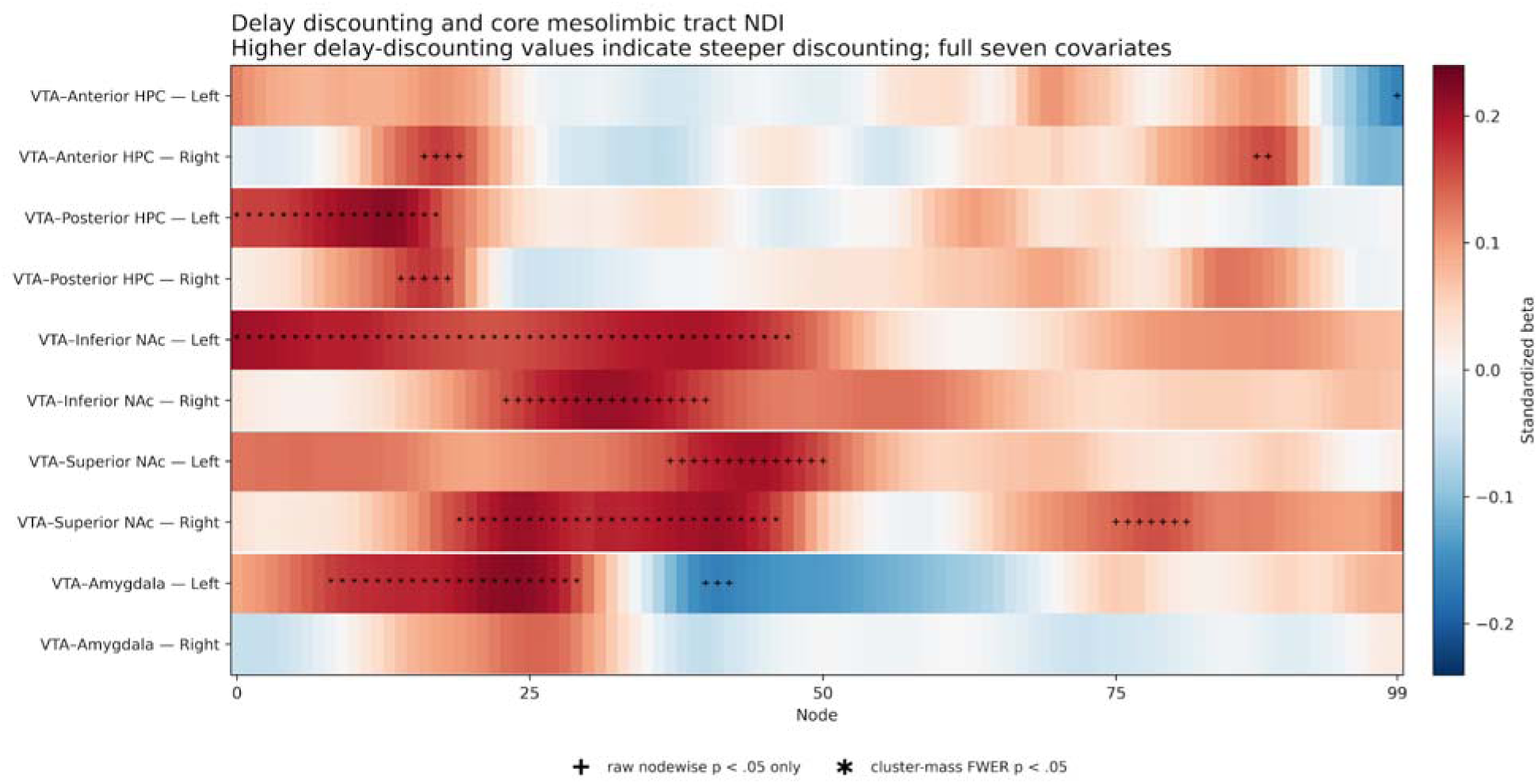
Nodewise NDI associations with delay discounting. Higher behavioral values indicate steeper discounting. Color denotes the standardized behavior coefficient at each of 100 unsmoothed nodes. Plus signs mark raw nodewise p < .05; asterisks mark nodes belonging to a within-profile cluster-mass-FWER-corrected cluster (p <.05). Correction was performed separately within each tract-by-hemisphere profile.

## Discussion

This study provides a pathway-resolved human atlas of mesolimbic circuitry and demonstrates its sensitivity to individual differences in motivated behavior. We reconstructed five VTA–centered pathways and two components of the HPC–NAc–VP–VTA downward arc. Inferior and superior VTA–nucleus accumbens (NAc) pathways occupied distinct accumbal territories, anterior and posterior VTA–hippocampus (HPC) pathways approached different long-axis and subfield territories, and the five VTA–centered pathways showed systematic midbrain topography. Neurite density covaried across mesolimbic pathways but not a comparison tract, and delay discounting was associated with localized microstructural variation in VTA–NAc, VTA–amygdala, and VTA–HPC pathways. Together, these findings provide a structural framework for testing how distinct but interacting mesolimbic pathways contribute to motivated learning, memory, and adaptive behavior.

### Topographic organization of human VTA–subcortical pathways

A central finding was that pathway identity could be recovered from anatomy at both ends of the reconstructed tracts. Distal endpoints differentiated inferior from superior VTA–NAc pathways and anterior from posterior VTA–HPC pathways. Within the VTA, all five direct mesolimbic pathways showed systematic spatial differences, and VTA endpoint coordinates classified pathway identity substantially above chance. Importantly, these distributions overlapped rather than forming discrete compartments. Thus, human VTA connectivity appears spatially organized without forming sharply segregated anatomical compartments.

This pattern closely parallels the heterogeneous but organized structure of dopamine signaling described in animal models. VTA dopamine neurons differ in molecular properties, physiology, afferent weighting, and projection targets, but they are not segregated into independent functional modules, and can multiplex reward, sensory, motor, and cognitive signals.^15,16,18,53^ At the same time, functional and projection-defined populations differ in the relative weighting of reward, sensory, and behavioral information.^19,20^ The present human anatomy is consistent with this organization. Pathway position may constrain the mixture of information available to downstream targets without imposing a fixed one-pathway, one-function correspondence. Mesolimbic organization may therefore be better characterized as a many-to-many mapping in which partially overlapping projection systems carry differently weighted mixtures of information.

The VTA-centered trajectories reconstructed here ascend through the broader territory historically associated with the medial forebrain bundle (MFB), itself a loosely organized collection of ascending and descending fibers rather than a single compact tract.^54,55^ Human tractography has proposed inferomedial and superolateral MFB routes,^56^ although the anatomical status of the superolateral route as a discrete fiber bundle remains debated.^57^ For this reason, the present atlas is organized around reproducible target-defined VTA trajectories rather than treating the MFB as a unitary pathway. Independent human anatomy supports these trajectories. Skandalakis et al.^25^ identified VTA-related connections with the NAc, amygdala, HPC, and entorhinal cortex using cadaveric microdissection and diffusion tractography. Their correspondence with MesoConnect provides an ex vivo-informed anatomical benchmark, while the present atlas resolves individual routes and quantifies their endpoint organization across participants.

### Motivated learning and salience circuit

The inferior and superior VTA–NAc pathways were separable both by trajectory and by their NAc endpoints. MacNiven et al.^24^ provided the most direct prior human comparison, resolving two midbrain–NAc trajectories using the anterior commissure as an anatomical landmark. Their inferior pathway coursed through lateral hypothalamic territory below the anterior commissure, whereas the superior pathway passed above the commissure through the anterior limb of the internal capsule and entered the NAc dorsally. We therefore used the anterior commissure to distinguish the same broad inferior and superior routes.

Our findings reproduce the inferior-superior route distinction described by MacNiven et al.^24^ but differ in midbrain organization. MacNiven et al. found the inferior NAc pathway more medial than the superior pathway, followed progressively laterally by caudate– and putamen–projecting pathways. They interpreted this gradient in relation to the ascending striatonigrostriatal spiral described in nonhuman primates, which links ventral, associative, and sensorimotor striatal territories through progressively more dorsolateral midbrain circuitry.^58,59^ However, MacNiven et al. examined a combined VTA/SN region, and the ascending spiral spans the broader dopaminergic midbrain and both ventral and dorsal striatum. By restricting endpoint analyses specifically to the VTA at higher anatomical resolution, we instead found the superior VTA–NAc pathway more medial than the inferior pathway.

This pattern comports with medial-to-medial and lateral-to-lateral organization described in tracing and projection-defined studies: medial VTA neurons preferentially innervate medial or dorsomedial NAc territories, whereas more lateral VTA populations preferentially innervate lateral accumbal fields.^12–14,29,60–66^. Modern projection-defined studies reinforce this organization: dopamine neurons projecting to the medial and dorsomedial NAc shell are concentrated in medial or ventromedial VTA territories, whereas lateral-shell-projecting neurons are shifted toward the lateral and dorsolateral VTA.^15–17,32,67–69^ The present findings parallel this organization: the superior VTA–NAc pathway occupied a more medial VTA position and approached a more medial and dorsal NAc territory, whereas the inferior pathway was positioned more laterally at both ends. Projection-defined populations remain partially intermingled, consistent with the substantial overlap observed within the VTA. The superior and inferior labels nevertheless describe trajectories around the anterior commissure and should not be treated as direct synonyms for dorsomedial versus lateral shell or for core versus shell.

Functional organization is similarly unlikely to follow a simple anatomical dichotomy. Projection-defined VTA populations convey different combinations of reward, aversion, and salience information depending on target and behavioral context. Lammel and colleagues linked lateral VTA to lateral NAc shell circuitry particularly strongly with reward and reinforcement while identifying aversion-related signaling in other projection-defined populations.^17,31,32^ In contrast, Cai et al.^30^ observed stronger reward-prediction-error-like signaling medially and stronger salience-like signaling laterally during fear extinction. Thus, the present anatomical separation may provide novel structural substrates for differently weighted motivational signals in humans, but it does not establish fixed reward prediction error or salience channels.

The VTA–amygdala pathway extends this motivated-learning framework beyond the NAc to affective salience and associative learning. Classical tracing identified substantial but overlapping amygdala-projecting populations within the VTA and adjoining substantia nigra.^70,71^ More recent work shows that midbrain-amygdala dopamine signaling contributes to salient sensory and associative learning and anxiety-related behavior,^34,35^ while amygdala lesions impair reinforcement learning from positive feedback.^33^ In humans, L-dopa increases temporal discounting, and individual susceptibility to this effect covaries with amygdala responses to reward proximity.^51^ Together, these findings suggest that VTA–amygdala circuitry contributes broadly to motivational significance and value-guided learning rather than carrying a single-valence signal, consistent with its inclusion alongside VTA–NAc pathways within the motivated learning and salience system examined here.

### The motivated memory circuit and hippocampal dopamine

The anterior and posterior VTA–HPC pathways approached markedly different hippocampal territories. Anterior endpoints were concentrated near the hippocampal head and were strongly weighted toward CA1 when they intersected participant-specific subfield labels. Posterior endpoints approached the hippocampal body and more frequently overlapped dentate gyrus labels. These findings provide human structural evidence that VTA–HPC connectivity is heterogeneous along the hippocampal long axis.

Comparative anatomy supports this organization. Tracing studies identify denser dopaminergic innervation of ventral hippocampal territories, particularly the subiculum and adjacent CA1, while also demonstrating projections to dorsal hippocampus.^36,38,39,72^ Human structural findings show a corresponding long-axis bias, with stronger VTA connectivity to anterior than posterior HPC.^43^ However, Skandalakis et al.^25^ further identified human VTA-related fibers extending toward dorsal dentate gyrus using tractography guided by cadaveric microdissection. Together, these findings support anatomically differentiated VTA–HPC connectivity along the hippocampal long axis rather than a homogeneous projection.

This anatomical separation is potentially important because anterior and posterior HPC support different representational functions. Posterior HPC is more strongly associated with fine-grained spatial and contextual representations, whereas anterior HPC is more strongly associated with coarse, generalized, and affective representations.^73–75^ VTA–Posterior HPC connectivity may therefore be particularly relevant to motivational modulation of precise contextual representations, whereas VTA–Anterior HPC connectivity may be more relevant to generalized or affectively significant memories. These functional assignments remain hypotheses, however, because diffusion tractography establishes anatomical trajectories rather than the computations carried by those pathways.

A substantial literature nevertheless establishes a broader role for VTA–HPC dopamine signaling in motivated memory. Dopamine D1/D5 receptor signaling facilitates late-phase CA1 long-term potentiation (LTP), novelty-dependent plasticity, and memory persistence^40,41,76,77^ Projection-specific experiments further demonstrate functional midbrain dopamine input to dorsal CA1 and show that VTA dopamine signaling can modulate contextual memory, hippocampal LTP, and contextual learning.^37,42^ Human fMRI studies similarly implicate VTA–HPC interactions in the selective encoding and consolidation of motivationally significant information. Reward anticipation engages the VTA and HPC before subsequently remembered events,^4,78^ and reward-related memory benefits are also associated with post-encoding interactions involving the VTA and anterior HPC.^79,80^ Greater midbrain–HPC tract density is also associated with enhanced memory for both monetary gains and losses, directly linking individual differences in structural connectivity to motivated memory.^22^ By separating anterior and posterior VTA–HPC pathways in vivo, the present atlas therefore provides an anatomically resolved framework for testing whether mesolimbic modulation differentially influences the distinct representations supported along the hippocampal long axis.

These pathways may also provide a useful framework for investigating neurodegenerative disease. Midbrain dopamine neuron loss contributes to hippocampal plasticity and memory deficits in animal models of Alzheimer’s disease, while stimulation of residual dopaminergic circuitry can partially rescue hippocampal synaptic deficits.^81,82^ Dopamine input to lateral entorhinal cortex also supports associative memory and is disrupted early in an amyloid precursor protein knock-in model.^83,84^ Although the present atlas cannot establish dopamine identity or an entorhinal continuation, separating anterior and posterior VTA–HPC trajectories provides more specific structural targets for future multimodal studies of dopamine-related memory vulnerability.

### Regulation of motivated behavior through the downward arc

The HPC–NAc–VP and VP–VTA reconstructions extend the atlas beyond direct VTA–centered pathways to a circuit proposed to regulate VTA dopamine responsivity. In the HPC–VTA loop framework, HPC activity engages the NAc and VP to regulate the number of spontaneously active VTA dopamine neurons. This population signal determines how many neurons are available to respond to later phasic input, thereby regulating the gain of the dopamine system.^3,26,44^ The downward arc therefore provides a mechanism through which contextual and novelty-related information can alter subsequent motivated responding.

Studies implicate components of this circuit in several forms of maladaptive behavior. Ventral HPC–NAc projections regulate susceptibility to depression-like behavior in animal models,^45^ hippocampal-striatal circuitry contributes to context-dependent drug seeking, and excessive hippocampal drive through ventral striatal and pallidal relays is proposed to produce dopamine dysregulation in psychosis.^46,47,85^ Human diffusion imaging likewise links greater HPC-ventral striatal structural connectivity to greater binge-drinking frequency.^48^ These findings reinforce the relevance of the downward arc to regulation of motivated behavior beyond memory alone.

### Coordinated microstructure across mesolimbic pathways

NDI covaried broadly across the mesolimbic pathways, with 20 of 21 pairwise relationships surviving correction in the primary analysis. The arcuate fasciculus did not show this covariance pattern, which remained largely intact after adjustment for whole-WM NDI. Mesolimbic microstructure therefore shows coordinated circuit-level variation rather than independent variation across pathways.

One possibility is that pathways participating in a common circuit undergo partially coordinated development or experience-dependent remodeling. Activity-dependent plasticity extends beyond synapses to axons and myelin, and repeated coactivation can alter white matter structure across functionally coupled systems.^86,87^ Mesolimbic white matter also shows behavioral plasticity: VTA oligodendrogenesis and myelin plasticity occur during normal opioid reward learning in mice.^88^ The present NDI correlations are compatible with coordinated circuit organization.

### Individual differences in adaptive choice

Delay discounting provided a proof of concept for applying the atlas to individual differences in motivated behavior. Steeper discounting was associated with localized NDI variation in left VTA–Inferior NAc, right VTA–Superior NAc, left VTA–Amygdala, and left VTA–Posterior HPC. This distributed localization is consistent with intertemporal choice depending on more than reward valuation alone. Choosing between immediate and delayed outcomes requires valuation and action selection, but also representation of future outcomes, retrieval of relevant experience, and sensitivity to motivational significance. The observed associations therefore identify candidate structural substrates through which these processes could contribute to individual differences in discounting.

The VTA–NAc findings converge with previous structural work. MacNiven et al.^24^ found that lower inferior VTA–NAc tract coherence predicted greater trait impulsivity across independent samples, distinguished individuals with stimulant use disorder, and was associated with delay discounting. Elliott et al.^23^ similarly linked greater midbrain-limbic striatal connectivity to impulsivity in adolescents and young adults with and without attention-deficit/hyperactivity disorder (ADHD), including delay discounting. Our finding that higher NDI was associated with steeper discounting therefore extends evidence that structural variation within VTA–NAc circuitry relates to impulsive choice, while localizing these associations continuously along both inferior and superior VTA–NAc trajectories. Despite differences in diffusion metrics, pathway definitions, and samples, these studies collectively support a relationship between VTA–NAc microstructure and individual differences in impulsive and value-guided behavior.

The VTA–amygdala and VTA–Posterior HPC findings broaden this pattern but warrant greater caution. The amygdala association is compatible with evidence that dopamine alters temporal discounting together with amygdala responses to reward proximity,^51^ while the posterior HPC finding is consistent with a potential contribution of contextual or future-oriented memory representations to intertemporal choice. These are exciting avenues for future investigations.

### Limitations and future directions

Several limitations constrain interpretation. Diffusion tractography estimates plausible white matter trajectories but cannot establish axonal direction, neurotransmitter identity, synaptic connectivity, or monosynaptic projection anatomy.^89,90^ (Jones et al., 2013; Maier-Hein et al., 2017). This is particularly important in the MFB and subthalamic region, where ascending, descending, and crossing systems occupy closely neighboring territory. Endpoint maps likewise indicate where reconstructed streamlines approach a region rather than histological terminal fields.

The results also depend on region definitions and tractography constraints. The VTA is small, internally heterogeneous, and adjacent to the substantia nigra and several major fiber systems. Although high-resolution 7 Tesla imaging and participant-native reconstruction improve anatomical precision, the resulting pathways cannot be assumed to consist exclusively of VTA-derived or dopaminergic axons. Similarly, hippocampal subfield analyses quantify spatial overlap between tractography endpoints and participant-specific segmentations rather than direct synaptic innervation of CA1 or dentate gyrus.

Finally, the present cohort consists of healthy young adults. Independent diffusion datasets, postmortem anatomy, and multimodal imaging will be important for testing generalizability and validating pathway geometry. Combining these atlases with dopamine positron emission tomography (PET) could test whether structural variation predicts transmitter-specific measures, while functional imaging could determine whether anatomically distinct pathways differ in task-related coupling. Longitudinal and clinical studies will be necessary to determine whether mesolimbic microstructure reflects stable predispositions, experience-dependent plasticity, or both.

## Conclusion

MesoConnect resolves the human mesolimbic system into distinct VTA–NAc, VTA–amygdala, VTA–HPC, and HPC–NAc–VP–VTA pathways with differentiated trajectory and endpoint organization. By integrating pathway geometry, endpoint topology, coordinated microstructure, and individual differences in behavior, this work advances a pathway-resolved account of human mesolimbic circuitry. The freely available MesoConnect atlas, reconstruction materials, and nodewise tractometry tutorial provide a reproducible framework for reconstructing these pathways in independent datasets and testing their relationships with cognition, behavior, and disease. MesoConnect atlas files, step-by-step documentation, and tutorials are available on the MesoConnect website (https://mesoconnect.vercel.app).

## Methods

### Participants and data source

Diffusion-weighted and structural magnetic resonance imaging data were obtained from the Human Connectome Project 7T Young Adult release.^91^ The initial candidate sample comprised 178 adults with 7T diffusion data. Three participants lacked required T1-weighted structural products and two had unusable diffusion-to-structural registration, yielding a tractography-eligible cohort of 173 adults (105 women, 68 men; age M = 29.47 years, SD = 3.32, range = 22–36 years). The HCP acquisition was approved by the Washington University institutional review board, all participants provided written informed consent, and the present study used deidentified data under the HCP data-use terms. Successful bilateral reconstruction varied by pathway because additional pathway-specific anatomical and quality-control criteria were applied (Tables 1, Supplementary Tables S1–S3). Each downstream analysis used the largest sample with all required pathway and analysis-specific products.

### Magnetic resonance imaging acquisition and HCP preprocessing

Diffusion magnetic resonance imaging was acquired on a 7T Siemens MAGNETOM scanner with a 70 mT/m gradient system and a 32-channel head coil. A multiband spin-echo echo-planar sequence was used with echo time = 71.2 ms, repetition time = 7,000 ms, flip angle = 90 degrees, refocusing flip angle = 180 degrees, field of view = 210 x 210 mm, matrix = 200 x 200, 132 slices, 1.05-mm isotropic voxels, multiband factor = 2, echo spacing = 0.82 ms, and bandwidth = 1,388 Hz/Px. The acquisition included b values of 1,000 and 2,000 s/mm2 and 65 diffusion-weighting directions.

Data were minimally preprocessed with the HCP diffusion pipeline.^92^ Structural images were corrected for gradient nonlinearity, readout distortion, and bias-field inhomogeneity. Diffusion images were intensity normalized to the mean b = 0 image and corrected for susceptibility-induced distortion, eddy-current distortion, participant motion, and gradient nonlinearity before alignment to structural space. The T1-weighted image therefore served as the anatomical reference for region-of-interest registration and native-space tractography.

### Fiber-orientation estimation

Preprocessed diffusion images were analyzed with MRtrix3 version 3.0.4.^93^ Voxelwise white-matter, gray-matter, and cerebrospinal-fluid response functions were estimated for each participant with dwi2response dhollander,^94^ and the participant-specific response functions were averaged with responsemean to obtain group white-matter, gray-matter, and cerebrospinal-fluid response functions. Multi-shell, multi-tissue constrained spherical deconvolution was then performed with dwi2fod msmt_csd.^95^ The resulting fiber-orientation distributions were bias corrected and intensity normalized across tissues and participants with mtnormalise.^96^

### Regions of interest and anatomical constraints

The pathway set comprised five bilateral VTA–centered tracts: VTA–Anterior HPC Tract, VTA–Posterior HPC Tract, Inferior VTA–NAc Tract, Superior VTA–NAc Tract, and VTA–Amygdala Tract. Additionally, there were two bilateral components of the HPC– striatopallidal–VTA regulatory arc, HPC–NAc–VP Tract, and VP–VTA Tract.

HPC, amygdala, and NAc masks were obtained from the Harvard-Oxford subcortical atlas. The VTA was defined with the probabilistic 7T human atlas of Trutti et al.,^97^ thresholded at 25% probability, and the VP was defined with the Pauli et al.^98^ atlas. Masks were transformed from Montreal Neurological Institute space to each participant’s T1-weighted space with Advanced Normalization Tools (ANTS^99^) and visually inspected for anatomical alignment.

Overlapping masks were refined before tracking. The hippocampal mask was subtracted from the amygdala where necessary. The VP was dilated by one voxel to reach the adjacent gray-white interface, and the dilated VP was subtracted from the limbic striatal mask. General exclusions included thalamus, cortical gray matter, cerebellum, brainstem inferior to the VTA, mammillary bodies, and red nucleus; the red nucleus was also subtracted from the VTA mask. Tract-specific inclusion and exclusion masks restricted reconstructions to the intended ipsilateral route. The body of the fornix was excluded during VTA–HPC and VTA–Amygdala tracking to prevent a large neighboring bundle from dominating those reconstructions. This exclusion was a tractography constraint rather than a claim that biological midbrain-medial temporal pathways uniformly avoid the fimbria-fornix system, for which comparative anatomy supports multiple routes.

### Probabilistic tractography and pathway segmentation

Probabilistic tractography was performed in participant-native T1w-aligned diffusion space with the iFOD2 probabilistic tracking algorithm implemented in MRtrix3 *tckgen*, using the intensity-normalized white-matter fiber-orientation distribution.^93,100^ Tracking was unidirectionally seeded from the pathway-specific seed mask, and the contralateral hemisphere was excluded. For every direct and downward-arc reconstruction, tracking continued until 2,500 accepted streamlines or 25 million seeding attempts had been reached, ensuring sufficient streamlines to generate a robust tractogram for atlas construction. No ACT or backtracking flags were used. The seed convention standardized reconstruction across participants and should not be interpreted as evidence of physiological direction. Final pathway-specific parameters and mask logic were selected after pilot reconstructions and then held fixed across participants (reported in Supplementary Tables S1 and S2).

The HPC–striato–pallidal–VTA downward arc was reconstructed as two separable pathway components. The HPC–NAc–VP pathway was seeded unidirectionally from the ipsilateral HPC and required ordered inclusion through the fornix, ipsilateral NAc, and ipsilateral VP; thus, the NAc served as an intermediate waypoint in the HPC-to-VP reconstruction. VP–VTA was reconstructed separately from an ipsilateral VP seed and ipsilateral VTA. Successful reconstruction of several mesolimbic pathways required substantially lowering the FOD threshold, which improved sensitivity but also increased susceptibility to spurious streamlines. We therefore did not retain attempts to force a single continuous HPC-to-VTA streamline. The additional waypoints imposed overly restrictive trajectory constraints, and, because the proposed circuit is polysynaptic, diffusion tractography cannot directly bridge its synaptic relays. Imposing continuous streamline geometry across the full arc under these conditions would risk selecting anatomically implausible false-positive trajectories. Accordingly, the two reconstructed components provide structural proxies for portions of the regulatory circuit rather than evidence for a single continuous monosynaptic tract.

### Streamline clustering and cleaning

When an initial tractogram contained a stray streamline cluster, QuickBundles clustering in Diffusion Imaging in Python (DIPY) was used to separate streamline clusters and retain the anatomically plausible component.^101,102^ Each retained tractogram was resampled to 100 equidistant nodes. At each node, the tract core was defined from the mean streamline coordinates, and streamlines exceeding 3 SD in Mahalanobis distance from the core were removed. Streamlines exceeding 2 SD from the mean tract length were also removed. Cleaning was repeated for at most five iterations and was followed by slice-by-slice visual quality control.

### Probabilistic atlas generation and atlas agreement

Cleaned tractograms were converted with MRtrix3 tckmap to voxel maps on each participant’s 1.05-mm T1-aligned grid. Subject maps were transformed to the 1-mm MNI152 grid with participant-specific ANTs transforms using nearest-neighbor interpolation, binarized, summed across valid participants, and divided by the number of valid maps for that pathway. Thresholded products were generated at 25%, 50%, and 75% participant overlap. The atlas-generation workflow is summarized in Figure 1.

For subject-to-atlas agreement, each 25%, 50%, and 75% group atlas was transformed from MNI space to the corresponding participant’s native tract grid using the participant-specific nonlinear warp and affine transform, nearest-neighbor interpolation, and the participant tract mask as the reference. The thresholded atlas and participant binary tract mask were compared using ordinary Dice, one-voxel-dilated Dice, the 95th-percentile symmetric Hausdorff distance (HD95), bundle overlap, and bundle overreach. Dilated Dice was calculated after one iteration of binary dilation of both masks, and HD95 was expressed in millimeters. Following standard tractography-evaluation conventions, bundle overlap was the intersection volume divided by participant-tract volume, whereas bundle overreach was atlas volume outside the participant tract divided by participant-tract volume.^90^ Thus, higher bundle overlap indicates greater participant-tract coverage and lower bundle overreach indicates less excess atlas territory. Tract-by-threshold medians were calculated across valid participants.

MRtrix3 tckstats was used to summarize the unweighted streamline-length distribution and raw accepted-streamline count for each valid participant-tract file. No streamline weights were supplied. Raw streamline count was treated as an algorithm-dependent tracking output rather than an estimate of axon number or biological connection strength.

### Endpoint topology

Streamline endpoints within the VTA and distal target regions were converted to endpoint-density maps on each participant’s 1.05-mm NODDI grid with MRtrix3 tckmap using the endpoint-only option. Density-weighted centers of mass were calculated in physical native-space coordinates and transformed to MNI space with participant-specific ANTs transforms. These summaries index where reconstructed streamlines approach a region of interest and should not be interpreted as histological synaptic terminal fields. Full command patterns, resampling rules, endpoint-availability criteria, and hippocampal-subfield analyses are provided in the Supplementary Methods.

Hippocampal endpoint topology was evaluated separately for bilateral VTA–Anterior HPC and VTA–Posterior HPC pathways using participant-specific FreeSurfer hippocampal-subfield segmentations resampled to the endpoint grid.^103^ Endpoint-density maps were generated on each participant’s native 1.05-mm NODDI grid using MRtrix3 3.0.4 with tckmap. We recorded the dominant overlapping subfield, hippocampal head versus body overlap, and endpoint position relative to MNI y = −21. Exact-overlap analyses were restricted to endpoint voxels intersecting a resampled subfield label, and a one-voxel dilation sensitivity analysis assessed whether nonoverlapping endpoint masks lay immediately adjacent to the subfield surface. Fisher exact tests compared anterior versus posterior center-of-mass classifications, Mann-Whitney U tests compared exact-overlap center-of-mass y coordinates, and a chi-square test compared dominant-subfield distributions.

NAc endpoint topology was compared between VTA–Inferior NAc and VTA–Superior NAc in the 173 participants with complete bilateral data. Left and right coordinates were collapsed by averaging absolute x, representing distance from the midline, and by averaging y and z. A paired three-coordinate multivariate test, equivalent to Hotelling’s T2 on the within-participant coordinate difference, tested separation between pathway classes.

VTA-wide topology was evaluated from VTA-adjacent endpoint centers of mass for the five direct pathway classes in the 156 participants with complete bilateral observations. ADistance from the midline (|x|), y, and z were compared with repeated-measures analyses, with Greenhouse-Geisser correction when appropriate, and paired multivariate contrasts were Holm corrected. Linear discriminant analysis used the three coordinates to predict pathway identity. Leave-one-subject-out validation excluded all five observations from one participant during model fitting and classified those observations together. Uncertainty in classification accuracy was estimated with 10,000 participant-level bootstrap samples.

### Neurite orientation dispersion and density imaging

The neurite orientation dispersion and density imaging model was fit with Accelerated Microstructure Imaging via Convex Optimization (AMICO) version 2.1.0,^104,105^ yielding neurite density index, orientation dispersion index, and free-water fraction maps. Primary analyses used volume-modulated NDI, a model-derived index sensitive to the intra-neurite signal fraction while reducing regional partial-volume bias.^106^ Nodewise tractometry followed the Automated Fiber Quantification framework in Python (pyAFQ), which represents tissue properties along a standardized trajectory for each bundle.^107,108^ Because the mesolimbic bundles had already been reconstructed and cleaned, we used the tract-profile stage rather than pyAFQ’s end-to-end bundle-recognition workflow. Streamlines were oriented within participant and pathway, resampled to 100 equal-arclength nodes, and NDI was sampled using core-weighted averaging. Whole-tract and nodewise NDI were retained for analysis. Full input requirements, interpolation and weighting rules, node-orientation procedures, and quality-control steps are provided in the Supplementary Methods. Whole-white-matter mean NDI was obtained from each participant’s FreeSurfer white-matter segmentation.^109^ The arcuate fasciculus was segmented with TractSeg, a standalone Python software package for automated white-matter tract segmentation,^110^ and served as a nonmesolimbic comparison bundle. It was selected to provide a spatially conservative control rather than an arbitrarily distant tract. Specifically, centroid distances were calculated in 1-mm MNI space between the bilateral 50%-overlap mask for each available mesolimbic pathway component and each of 41 bilateral or midline TractSeg bundles thresholded at 50% probability. The arcuate fasciculus was selected as a spatially conservative nonmesolimbic control because we sought a major long-range association pathway that was spatially proximal to, but anatomically distinct from, the mesolimbic system. Among the 41 TractSeg bundles evaluated, it had the smallest mean centroid distance across the eight mesolimbic component masks (M = 18.36 mm, range = 14.90–20.39 mm), while showing no overlap with any mesolimbic 50% atlas mask (Dice = 0; minimum voxel distance = 2.00–19.85 mm), supporting its use as an anatomically nearby but nonmesolimbic control.

The primary circuit-covariation analysis included bilateral-mean whole-tract NDI for HPC–NAc–VP, VP–VTA, and the five direct VTA–centered pathways, with arcuate fasciculus NDI as a comparison. Pairwise partial Pearson correlations controlled for sex, age, intracranial volume, and handedness. Sensitivity analysis further investigated this relation controlling for whole-white-matter NDI. Observations more than 3 SD from either raw tract variable were excluded once before covariate residualization. Benjamini-Hochberg false-discovery-rate correction was applied to two prespecified families within each specification: the 21 mesolimbic-mesolimbic pairs and the seven arcuate-versus-mesolimbic comparisons.

Behavioral associations of the VTA–centered (i.e., “upward-arc”) mesolimbic tracts were investigated using delay discounting, a task hypothesized to broadly engage the circuit.^49–51,111^ Delay discounting was quantified as the area under the empirical discounting curve,^112^ and scores were sign-reversed and standardized so that higher values indicated steeper discounting, or a stronger preference for smaller immediate rewards over larger delayed rewards. Positive regression coefficients therefore indicate higher NDI with greater delay discounting (impulsivity). At every node of each tract and hemisphere, NDI was modeled as a function of standardized discounting with sex, age, intracranial volume, handedness, raw streamline count, and mean streamline length included as covariates. The nodewise delay-discounting analysis tested all five prespecified upward-arc pathways bilaterally: the inferior VTA–NAc tract, superior VTA–NAc tract, VTA–amygdala, VTA–anterior HPC, and VTA–posterior HPC. The two downward-arc pathways, HPC–NAc–VP and VP–VTA, were prespecified atlas targets but were not included in this nodewise analysis.

Inference used 10,000 response-side Freedman-Lane permutations.^113^ NDI profiles and behavior were residualized on the nuisance design. At each permutation, complete reduced-model NDI residual rows were permuted across participants, projected off nuisance space again, and tested against the fixed covariate-adjusted behavior. The same row permutation was used across all 100 nodes, preserving within-profile node covariance. Traditional cluster-mass correction used a two-sided nodewise cluster-forming threshold of raw p < .05.^114^ Positive and negative contiguous clusters were formed separately, cluster mass was defined as the sum of absolute node t values, and the maximum cluster mass across the 100-node profile was retained for each permutation.

## Data availability

Human Connectome Project (HCP) source imaging data are available through ConnectomeDB, subject to HCP data-use terms. Derived MesoConnect probability atlases at unthresholded, 25%, 50%, and 75% participant-overlap levels, region definitions, endpoint products, tractometry profiles, and quality-control materials are openly available through the MesoConnect GitHub repository (https://github.com/blelliott23/MesoConnect). Documentation, download instructions, and tutorials for applying the atlas are available on the MesoConnect website (https://mesoconnect.vercel.app).

## Code availability

Custom code used for atlas construction, endpoint analyses, subject-to-atlas agreement, circuit-level neurite density index analyses, tractometry, and permutation-based nodewise inference is openly available through the MesoConnect GitHub repository (https://github.com/blelliott23/MesoConnect).

## Acknowledgments

This work used data provided by the Human Connectome Project, Washington University–Minnesota Consortium. The authors thank the Human Connectome Project participants and investigators.

## Author Contributions

B.L.E. and R.M. conceived the study and developed and implemented the tractography and analysis framework. B.L.E. drafted the manuscript. R.M., L.J.H., J.K.L., N.D.V., I.R.O. and V.P.M. contributed to study framing, interpretation and critical revision of the manuscript. All authors reviewed and approved the manuscript.

