## Supplemental Methods and Tables for "Mapping Mesolimbic Dopamine Projections to Subcortical Circuits Underlying Adaptive Behavior: A Human 7T Diffusion Tractography Atlas" (from another version of this preprint: version 1, 2026-09-28)

**Supplementary Methods**

The main Methods describe the analytical rationale and primary statistical models. The details below document the implementation choices needed to reproduce the endpoint and nodewise analyses without repeating the main text. Pathway-specific tracking settings and mask logic are summarized in Supplementary Tables S1 and S2, and atlas-agreement definitions and results are reported in Supplementary Table S3.

**Endpoint-density maps and coordinate handling**

Final endpoint-density maps were generated from each cleaned tractogram with the MRtrix3 command pattern tckmap <cleaned_tract.tck> <endpoint_density.nii.gz> -ends_only -template <fit_NDI.nii.gz>. The template placed each map on the participant’s native 1.05-mm NODDI grid. For every nonzero endpoint voxel, physical coordinates were derived from the NIfTI affine, and the endpoint center of mass was calculated as the endpoint-density-weighted mean coordinate. Participant-specific ANTs transforms were used to map endpoint coordinates and masks to the 1-mm MNI152 reference. Binary masks and anatomical labels were resampled with nearest-neighbor interpolation; continuous scalar images were sampled with trilinear interpolation.

**Hippocampal subfield and long-axis analyses**

T1-weighted structural MRI images were processed using the full FreeSurfer pipeline (version 8.0.0) for automated cortical and subcortical segmentation. Processing included the standard recon-all workflow, encompassing intensity normalization, skull stripping, surface reconstruction, tissue segmentation, and volumetric segmentation. Hippocampal subfields were additionally segmented using FreeSurfer’s automated hippocampal subfield segmentation procedure, providing volumetric estimates for individual hippocampal subregions (CA1, CA3, CA4, DG, Subiculum, Presubiculum, and Parasubiculum). Note that due to difficulty in resolving anatomical boundaries between CA2 and CA3, CA2 is subsumed into CA3. Subfields were concatenated across hemispheres yielding bilateral subfield ROIs that were visually inspected prior to subsequent endpoint topography analyses.

Participant-specific FreeSurfer hippocampal-subfield segmentations were resampled directly to the native endpoint grid with NiBabel resample_from_to using nearest-neighbor interpolation (order = 0). Exact overlap required a nonzero endpoint voxel to intersect a resampled subfield label. For each participant, pathway, and hemisphere, the dominant subfield was the label containing the largest number of exact-overlap endpoint voxels; counts for all intersected labels were retained. Head-versus-body overlap was summarized from the corresponding FreeSurfer labels, and anterior-posterior class was defined from the MNI-space endpoint center of mass relative to y = −21 mm.

Endpoint maps without exact label overlap were evaluated in a separate adjacency analysis by dilating the participant-specific subfield labels by one voxel on the native endpoint grid. This sensitivity analysis tested whether an endpoint immediately abutted a subfield boundary; it did not replace the undilated labels or alter the primary exact-overlap classifications. Fisher’s exact tests compared anterior-posterior classes, two-sided Mann–Whitney U tests compared exact-overlap MNI y coordinates, and chi-square tests compared dominant-subfield distributions.

**VTA endpoint topology and pathway classification**

For each participant and direct pathway, bilateral VTA-adjacent endpoint coordinates were summarized as mean absolute x (distance from the midline), mean y, and mean z. Analyses used the 156 participants with complete observations for all five pathways. One-way repeated-measures models were implemented with statsmodels AnovaRM; Greenhouse–Geisser epsilon, adjusted degrees of freedom, and adjusted p values were calculated from the within-participant covariance matrix. Paired three-coordinate Hotelling tests were adjusted across the 10 pathway pairs using Holm’s method. Pathway identity was predicted from absolute x, y, and z with scikit-learn LinearDiscriminantAnalysis. Each leave-one-subject-out fold excluded all five pathway observations from one participant during fitting and classified those five observations together. The 95% confidence interval for accuracy was estimated from 10,000 bootstrap resamples of participant-level accuracies. Complete outputs are reported in Supplementary Table S5.

**AFQ tract profiles and MesoConnect tutorial**

Automated Fiber Quantification represents a diffusion-derived scalar as an ordered profile along a bundle rather than as a single tract average (Yeatman et al., 2012; Kruper et al., 2021). Because the MesoConnect pathways were reconstructed and cleaned before tractometry, we reproduced the AFQ profile stage with DIPY/pyAFQ-compatible operations rather than using pyAFQ to recognize the bundles. Inputs were a cleaned native-space TCK file and an NDI NIfTI image on the corresponding native grid. Within each participant and pathway, an infinite-threshold QuickBundles cluster supplied one reference centroid; orient_by_streamline aligned all streamlines to that centroid, and set_number_of_points resampled every streamline to 100 equal-arclength nodes.

NDI was sampled at every streamline-node coordinate using trilinear interpolation. At each node, DIPY gaussian_weights downweighted streamlines according to their Mahalanobis distance from the bundle core and normalized the weights to sum to one. Despite the function name, these are inverse-Mahalanobis weights rather than an exponential Gaussian kernel. The node value was the weighted mean across streamlines. Each retained profile contained 100 ordered, finite node values. Orientation was standardized within each pathway profile; the same node number was not assumed to represent an equivalent anatomical location across different pathway classes.

The MesoConnect tutorial accepts cleaned pathway tractograms and scalar maps, verifies coordinate compatibility, and produces a long-format table containing participant, pathway, hemisphere, node, and scalar value. It also demonstrates the covariate-adjusted nodewise model and participant-level permutation structure used here and exports orientation and profile plots for quality control. The tutorial is intended to apply the released pathway definitions to independent data; it does not treat streamline count as axon number or nodewise NDI as a direct measure of dopamine signaling.

**Supplementary Table S1**

*Core Pathway-Specific Tractography Parameters*

| **Pathway** | **Seed and required inclusion order** | **FOD cutoff** | **Length, mm** | **Angle** | **Step, mm** |
| --- | --- | --- | --- | --- | --- |
| Inferior VTA–NAc tract | VTA → ipsilateral NAc | .03 | 8–35 | 7° | .25 |
| Superior VTA–NAc tract | VTA → ipsilateral NAc | .03 | 8–35 | Default | Default |
| VTA–amygdala | VTA → ipsilateral amygdala | .08 | 27–40 | Default | Default |
| VTA–posterior HPC | VTA → ipsilateral pHPC | .06 | 35–65 | Default | Default |
| VTA–anterior/posterior HPC | VTA→ ipsilateral aHPC | .06 | 35–65 | Default | Default |
| HPC–NAc–VP | HPC → fornix → NAc → VP | .04 | Minimum 5; no maximum | Default | Default |
| VP–VTA | VP → VTA | .03 | 15–25 | 15° | Default |

*Note.* All rows used the normalized white-matter FOD, unidirectional seeding, 2,500 accepted streamlines, a ceiling of 25 million seeding attempts, and the -stop option.

**Supplementary Table S2**

*Tract-Specific Mask Logic Confirmed in the Supplied Commands*

| **Pathway** | **Key confirmed exclusion logic** |
| --- | --- |
| Inferior VTA–NAc tract | Ipsilateral amygdala and HPC, contralateral NAc and hemisphere, eroded (1 voxel) thalamus, cortex/cerebellum, inferior brainstem, red nucleus, and anterior-superior anterior-commissure mask. |
| Superior VTA–NAc tract | Ipsilateral amygdala and HPC, contralateral NAc and hemisphere, eroded (1 voxel) thalamus, cortex/cerebellum, inferior brainstem, red nucleus, and anterior-inferior anterior-commissure mask. |
| VTA–posterior HPC | Ipsilateral VP, bilateral NAc, striatum, eroded (1 voxel) thalamus, cortex/cerebellum, inferior brainstem, amygdala, red nucleus, fornix, ipsilateral optic tract/nerves, and contralateral hemisphere. |
| VTA-anterior HPC | Ipsilateral VP, bilateral NAc, striatum, eroded (1 voxel) thalamus, cortex/cerebellum, inferior brainstem, red nucleus, fornix, ipsilateral optic tract/nerves, and contralateral hemisphere. |
| VTA–amygdala | Ipsilateral VP, NAc, HPC, optic tract/nerves, striatum, eroded (1 voxel) thalamus, cortex/cerebellum, inferior brainstem, red nucleus, fornix, and contralateral hemisphere. |
| HPC–NAc–VP | Cortex/cerebellum,, ipsilateral amygdala, and contralateral hemisphere. |
| VP–VTA | Fornix, eroded (1 voxel) thalamus, cortex/cerebellum, inferior brainstem, ipsilateral HPC and amygdala, and contralateral hemisphere. |

**Supplementary Table S3**

*Median Subject-to-Atlas Agreement by Tract and Atlas Threshold*

| **Tract** | **Threshold** | ***n*** | **Median Dice** | **Median dilated Dice** | **Median HD95 (mm)** | **Median BOL** | **Median BOR** |
| --- | --- | --- | --- | --- | --- | --- | --- |
| Left inferior VTA–NAc tract | 25% | 173 | 0.748 | 0.805 | 1.485 | 0.872 | 0.475 |
| Left inferior VTA–NAc tract | 50% | 173 | 0.737 | 0.800 | 1.485 | 0.686 | 0.173 |
| Left inferior VTA–NAc tract | 75% | 173 | 0.582 | 0.675 | 2.100 | 0.433 | 0.038 |
| Right inferior VTA–NAc tract | 25% | 173 | 0.752 | 0.809 | 1.485 | 0.849 | 0.423 |
| Right inferior VTA–NAc tract | 50% | 173 | 0.730 | 0.795 | 1.485 | 0.666 | 0.139 |
| Right inferior VTA–NAc tract | 75% | 173 | 0.578 | 0.673 | 2.100 | 0.426 | 0.026 |
| Left superior VTA–NAc tract | 25% | 173 | 0.767 | 0.825 | 1.485 | 0.896 | 0.432 |
| Left superior VTA–NAc tract | 50% | 173 | 0.765 | 0.822 | 1.485 | 0.731 | 0.146 |
| Left superior VTA–NAc tract | 75% | 173 | 0.657 | 0.741 | 1.485 | 0.516 | 0.035 |
| Right superior VTA–NAc tract | 25% | 173 | 0.770 | 0.825 | 1.485 | 0.893 | 0.398 |
| Right superior VTA–NAc tract | 50% | 173 | 0.768 | 0.826 | 1.420 | 0.705 | 0.137 |
| Right superior VTA–NAc tract | 75% | 173 | 0.626 | 0.724 | 1.819 | 0.472 | 0.030 |
| Left VTA–amygdala | 25% | 168 | 0.679 | 0.754 | 2.100 | 0.810 | 0.523 |
| Left VTA–amygdala | 50% | 168 | 0.646 | 0.724 | 2.286 | 0.574 | 0.168 |
| Left VTA–amygdala | 75% | 168 | 0.465 | 0.576 | 3.320 | 0.320 | 0.032 |
| Right VTA–amygdala | 25% | 168 | 0.706 | 0.772 | 1.819 | 0.821 | 0.465 |
| Right VTA–amygdala | 50% | 168 | 0.675 | 0.751 | 1.819 | 0.602 | 0.161 |
| Right VTA–amygdala | 75% | 168 | 0.507 | 0.613 | 2.572 | 0.360 | 0.030 |
| Left VTA–anterior HPC | 25% | 167 | 0.706 | 0.779 | 1.819 | 0.828 | 0.532 |
| Left VTA–anterior HPC | 50% | 167 | 0.693 | 0.767 | 1.819 | 0.617 | 0.184 |
| Left VTA–anterior HPC | 75% | 167 | 0.513 | 0.623 | 2.348 | 0.364 | 0.044 |
| Right VTA–anterior HPC | 25% | 167 | 0.670 | 0.762 | 1.819 | 0.802 | 0.546 |
| Right VTA–anterior HPC | 50% | 167 | 0.644 | 0.735 | 1.819 | 0.580 | 0.177 |
| Right VTA–anterior HPC | 75% | 167 | 0.452 | 0.578 | 2.572 | 0.308 | 0.045 |
| Left VTA–posterior HPC | 25% | 168 | 0.726 | 0.787 | 1.485 | 0.855 | 0.458 |
| Left VTA–posterior HPC | 50% | 168 | 0.674 | 0.746 | 2.100 | 0.614 | 0.141 |
| Left VTA–posterior HPC | 75% | 168 | 0.491 | 0.602 | 3.210 | 0.342 | 0.022 |
| Right VTA–posterior HPC | 25% | 168 | 0.710 | 0.776 | 1.882 | 0.845 | 0.437 |
| Right VTA–posterior HPC | 50% | 168 | 0.647 | 0.730 | 2.100 | 0.600 | 0.132 |
| Right VTA–posterior HPC | 75% | 168 | 0.478 | 0.585 | 3.150 | 0.320 | 0.028 |
| Left HPC–NAc–VP | 25% | 166 | 0.669 | 0.745 | 2.432 | 0.810 | 0.649 |
| Left HPC–NAc–VP | 50% | 166 | 0.661 | 0.730 | 2.572 | 0.597 | 0.193 |
| Left HPC–NAc–VP | 75% | 166 | 0.461 | 0.561 | 4.064 | 0.308 | 0.028 |
| Right HPC–NAc–VP | 25% | 166 | 0.662 | 0.746 | 2.348 | 0.815 | 0.630 |
| Right HPC–NAc–VP | 50% | 166 | 0.627 | 0.708 | 2.572 | 0.565 | 0.163 |
| Right HPC–NAc–VP | 75% | 166 | 0.403 | 0.517 | 4.200 | 0.261 | 0.019 |
| Left VP–VTA | 25% | 169 | 0.684 | 0.754 | 2.100 | 0.836 | 0.517 |
| Left VP–VTA | 50% | 170 | 0.654 | 0.735 | 2.100 | 0.594 | 0.173 |
| Left VP–VTA | 75% | 170 | 0.427 | 0.548 | 3.141 | 0.282 | 0.026 |
| Right VP–VTA | 25% | 170 | 0.692 | 0.761 | 2.100 | 0.842 | 0.524 |
| Right VP–VTA | 50% | 168 | 0.669 | 0.745 | 2.100 | 0.626 | 0.174 |
| Right VP–VTA | 75% | 170 | 0.471 | 0.589 | 2.988 | 0.328 | 0.030 |

*Note.* Values are medians across valid participants. Dice and dilated Dice range from 0 to 1, with higher values indicating greater agreement. Dilated Dice was calculated after one-voxel binary dilation of both masks. HD95 = 95th-percentile symmetric Hausdorff distance; lower values indicate closer surface agreement. BOL = bundle overlap, the fraction of participant-tract volume covered by the atlas; BOR = bundle overreach, atlas-only volume divided by participant-tract volume. Higher BOL and lower BOR are favorable. HPC = hippocampus; NAc = nucleus accumbens; VP = ventral pallidum; VTA = ventral tegmental area.

**Supplementary Table S4**

*Exact Statistics Corresponding to Figure 6 Circuit Level Covariation in Bilateral Mean Whole Tract Neurite Density Index*

| **Pathway pair** | **Sex-, age-, ICV-, and handedness-adjusted** | | | | **Sex-, age-, ICV-, handedness-, and whole-WM NDI-adjusted** | | | |
| --- | --- | --- | --- | --- | --- | --- | --- | --- |
|  | ***n*** | ***Partial r*** | ***p*** | ***q*** | ***n*** | ***Partial r*** | ***p*** | ***q*** |
| ***Mesolimbic pathway pairs*** | | | | | | | | |
| HPC–NAc–VP with VTA–anterior HPC | 158 | .2421 | .002568 | .002997 | 158 | .2150 | .007806 | .009107 |
| HPC–NAc–VP with VTA–posterior HPC | 161 | .2625 | .000931 | .001150 | 161 | .2147 | .007312 | .009033 |
| HPC–NAc–VP with VTA–amygdala | 160 | .2050 | .010522 | .011630 | 160 | .1758 | .029149 | .030606 |
| HPC–NAc–VP with Inferior VTA–NAc tract | 164 | .1535 | .053359 | .053359 | 164 | .1583 | .047002 | .047002 |
| HPC–NAc–VP with Superior VTA–NAc tract | 162 | .2853 | .000292 | .000383 | 162 | .2414 | .002401 | .003151 |
| HPC–NAc–VP with VP–VTA | 158 | .2007 | .012870 | .013513 | 158 | .1809 | .025701 | .028407 |
| VTA–anterior HPC with VTA–posterior HPC | 162 | .6963 | 4.37400e−24 | 3.06180e−23 | 162 | .6861 | 4.88075e−23 | 3.41653e−22 |
| VTA–anterior HPC with VTA–amygdala | 162 | .7829 | 9.19230e−34 | 1.93038e−32 | 162 | .7767 | 9.87114e−33 | 2.07294e−31 |
| VTA–anterior HPC with Inferior VTA–NAc tract | 166 | .3945 | 2.24198e−7 | 3.92346e−7 | 166 | .3984 | 1.80973e−7 | 3.80044e−7 |
| VTA–anterior HPC with Superior VTA–NAc tract | 165 | .3992 | 1.71103e−7 | 3.26651e−7 | 165 | .3731 | 1.27628e−6 | 2.06169e−6 |
| VTA–anterior HPC with VP–VTA | 160 | .3024 | .000131 | .000183 | 160 | .2922 | .000236 | .000330 |
| VTA–posterior HPC with VTA–amygdala | 163 | .7068 | 3.17182e−25 | 3.33041e−24 | 163 | .6996 | 2.16873e−24 | 2.27716e−23 |
| VTA–posterior HPC with Inferior VTA–NAc tract | 168 | .4395 | 4.36551e−9 | 1.30965e−8 | 168 | .4552 | 1.15846e−9 | 4.86555e−9 |
| VTA–posterior HPC with Superior VTA–NAc tract | 167 | .4450 | 2.98268e−9 | 1.25272e−8 | 167 | .4015 | 1.30106e−7 | 3.03581e−7 |
| VTA–posterior HPC with VP–VTA | 162 | .4487 | 3.79222e−9 | 1.30965e−8 | 162 | .4353 | 1.35451e−8 | 4.06354e−8 |
| VTA–amygdala with Inferior VTA–NAc tract | 168 | .4200 | 2.38681e−8 | 5.56921e−8 | 168 | .4239 | 1.90674e−8 | 5.00520e−8 |
| VTA–amygdala with Superior VTA–NAc tract | 167 | .3166 | 4.04103e−5 | 6.06154e−5 | 167 | .2920 | .000171 | .000257 |
| VTA–amygdala with VP–VTA | 163 | .3817 | 7.47068e−7 | 1.20680e−6 | 163 | .3767 | 1.15490e−6 | 2.02108e−6 |
| Inferior VTA–NAc tract with Superior VTA–NAc tract | 172 | .4285 | 7.60342e−9 | 1.99590e−8 | 172 | .4388 | 3.35404e−9 | 1.17391e−8 |
| Inferior VTA–NAc tract with VP–VTA | 167 | .5840 | 3.44183e−16 | 1.80696e−15 | 167 | .5868 | 2.83816e−16 | 1.49003e−15 |
| Superior VTA–NAc tract with VP–VTA | 166 | .3986 | 1.62929e−7 | 3.26651e−7 | 166 | .3860 | 4.64009e−7 | 8.85835e−7 |
| ***Arcuate fasciculus comparisons*** | | | | | | | | |
| HPC–NAc–VP with Arcuate fasciculus | 164 | .1679 | .034367 | .099685 | 164 | -.1308 | .101292 | .236348 |
| VTA–anterior HPC with Arcuate fasciculus | 166 | .0509 | .521501 | .608418 | 166 | -.2715 | .000516 | .003615 |
| VTA–posterior HPC with Arcuate fasciculus | 168 | .1589 | .042722 | .099685 | 168 | -.1890 | .016005 | .056016 |
| VTA–amygdala with Arcuate fasciculus | 168 | .0894 | .256169 | .358636 | 168 | -.1050 | .183792 | .321635 |
| Inferior VTA–NAc tract with Arcuate fasciculus | 173 | -.0128 | .869680 | .869680 | 173 | -.0337 | .665593 | .776525 |
| Superior VTA–NAc tract with Arcuate fasciculus | 172 | .2025 | .008681 | .060766 | 172 | .0079 | .919612 | .919612 |
| VP–VTA with Arcuate fasciculus | 167 | .0974 | .217387 | .358636 | 167 | .0372 | .639836 | .776525 |

*Note.* The sex-, age-, intracranial-volume (ICV)-, and handedness-adjusted model includes those four covariates. The whole-white-matter NDI-adjusted model includes the same covariates plus whole-white-matter neurite density index (NDI). Values are bilateral mean whole-tract NDI partial Pearson correlations. Two-sided p values are from conventional participant-level partial-correlation tests. Benjamini–Hochberg q values were calculated separately for the 21 mesolimbic pathway-pair tests and the seven arcuate-fasciculus comparisons within each model. HPC = hippocampus; NAc = nucleus accumbens; VP = ventral pallidum; VTA = ventral tegmental area.

**Supplementary Table S5. Panel A. Omnibus VTA Endpoint-Coordinate Tests**

| **Axis** | **F** | **df1** | **df2** | **Exact p** | **GG ε** | **Generalized η²** |
| --- | --- | --- | --- | --- | --- | --- |
| Distance from midline, \|x\| | 517.55 | 2.78 | 430.37 | 5.73e−137 | .694 | .489 |
| Anterior–posterior, y | 298.82 | 2.84 | 439.56 | 2.62e−102 | .709 | .561 |
| Superior–inferior, z | 93.32 | 3.13 | 485.38 | 3.06e−49 | .783 | .226 |

***Note.*** N = 156. Greenhouse–Geisser-adjusted degrees of freedom and exact p values are shown.

**Supplementary Table S5. Panel B. Paired Three-Coordinate Pathway Contrasts**

| **Pathway 1** | **Pathway 2** | **n** | **T²** | **F** | **df** | **Exact p** | **Holm p** |
| --- | --- | --- | --- | --- | --- | --- | --- |
| Inferior VTA–NAc tract | Superior VTA–NAc tract | 156 | 1628.92 | 535.97 | 3, 153 | 6.41e−81 | 4.49e−80 |
| Inferior VTA–NAc tract | VTA–posterior HPC | 156 | 400.53 | 131.79 | 3, 153 | 3.29e−42 | 9.86e−42 |
| Inferior VTA–NAc tract | VTA–amygdala | 156 | 617.64 | 203.22 | 3, 153 | 3.79e−53 | 1.90e−52 |
| Inferior VTA–NAc tract | VTA–anterior HPC | 156 | 406.16 | 133.64 | 3, 153 | 1.52e−42 | 6.09e−42 |
| Superior VTA–NAc tract | VTA–posterior HPC | 156 | 2979.85 | 980.47 | 3, 153 | 1.22e−99 | 1.22e−98 |
| Superior VTA–NAc tract | VTA–amygdala | 156 | 1875.07 | 616.96 | 3, 153 | 3.28e−85 | 2.62e−84 |
| Superior VTA–NAc tract | VTA–anterior HPC | 156 | 2870.04 | 944.34 | 3, 153 | 1.86e−98 | 1.67e−97 |
| VTA–posterior HPC | VTA–amygdala | 156 | 365.40 | 120.23 | 3, 153 | 4.80e−40 | 9.60e−40 |
| VTA–posterior HPC | VTA–anterior HPC | 156 | 116.21 | 38.24 | 3, 153 | 1.69e−18 | 1.69e−18 |
| VTA–amygdala | VTA–anterior HPC | 156 | 621.10 | 204.36 | 3, 153 | 2.69e−53 | 1.62e−52 |

***Note.*** Each contrast jointly tested |x|, y, and z with a paired Hotelling test. Holm adjustment covered all 10 contrasts.

**Supplementary Table S5. Panel C. Leave-One-Subject-Out Classification Sensitivity**

| **Pathway** | **Correct** | **Total** | **Sensitivity** | **Descriptive 95% CI** |
| --- | --- | --- | --- | --- |
| Inferior VTA–NAc tract | 115 | 156 | 73.7% | 66.3–80.0% |
| Superior VTA–NAc tract | 153 | 156 | 98.1% | 94.5–99.3% |
| VTA–posterior HPC | 91 | 156 | 58.3% | 50.5–65.8% |
| VTA–amygdala | 111 | 156 | 71.2% | 63.6–77.7% |
| VTA–anterior HPC | 91 | 156 | 58.3% | 50.5–65.8% |

***Note.*** Overall and balanced accuracy were 71.92% (561/780); the 95% participant-level bootstrap CI for overall accuracy was 69.23%–74.62%.

**Supplementary Table S6. Panel A. Endpoint Availability, Exact Overlap, and One-Voxel Adjacency**

| Pathway | Side | Available, n | Exact overlap, n (%) | Additional contact after one-voxel dilation, n | Final contact, n (%) |
| --- | --- | --- | --- | --- | --- |
| VTA–anterior HPC | Left | 166 | 80 (48.2) | 85 | 165 (99.4) |
| VTA–anterior HPC | Right | 166 | 92 (55.4) | 74 | 166 (100.0) |
| VTA–posterior HPC | Left | 168 | 160 (95.2) | 8 | 168 (100.0) |
| VTA–posterior HPC | Right | 168 | 157 (93.5) | 11 | 168 (100.0) |

***Note.*** Final contact combines exact overlap with additional contact after one-voxel dilation; one left VTA–anterior HPC endpoint remained without contact. HPC = hippocampus; VTA = ventral tegmental area.

**Supplementary Table S6. Panel B. Dominant Subfield by Endpoint Overlap**

| **Pathway** | **Side** | **n** | **CA1** | **CA3** | **CA4** | **DG** | **Presub.** | **Subic.** | **Parasub.** |
| --- | --- | --- | --- | --- | --- | --- | --- | --- | --- |
| VTA–anterior HPC | Left | 165 | 164 | 0 | 0 | 0 | 0 | 1 | 0 |
| VTA–anterior HPC | Right | 166 | 164 | 2 | 0 | 0 | 0 | 0 | 0 |
| VTA–posterior HPC | Left | 168 | 0 | 21 | 6 | 122 | 15 | 4 | 0 |
| VTA–posterior HPC | Right | 168 | 1 | 28 | 6 | 109 | 15 | 9 | 0 |

***Note.*** Dominant subfield was the label containing the largest number of endpoint voxels. DG = dentate gyrus; HPC = hippocampus; Parasub. = parasubiculum; Presub. = presubiculum; Subic. = subiculum; VTA = ventral tegmental area.

**Supplementary Table S6. Panel C. Long-Axis Position and Head–Body Distribution**

| **Pathway** | **Side** | **n** | **Head-dominant maps** | **Body-dominant maps** | **Ties** | **Head contact voxels, %** | **Body contact voxels, %** |
| --- | --- | --- | --- | --- | --- | --- | --- |
| VTA–anterior HPC | Left | 165 | 165 | 0 | 0 | 100.0 | 0.0 |
| VTA–anterior HPC | Right | 166 | 166 | 0 | 0 | 100.0 | 0.0 |
| VTA–posterior HPC | Left | 168 | 0 | 168 | 0 | 3.8 | 96.2 |
| VTA–posterior HPC | Right | 168 | 2 | 165 | 1 | 3.9 | 96.1 |

*Note****.*** The prespecified center-of-mass boundary

was MNI y = −21.

**Supplementary Table S6. Panel D. Route-Difference Tests**

| **Sample** | **Test** | **Outcome** | **Anterior n** | **Posterior n** | **Statistic** | **df** | **p** |
| --- | --- | --- | --- | --- | --- | --- | --- |
| Left | Fisher exact | Head- vs body-dominant map | 165 | 168 | ∞ | — | 1.32578 × 10⁻⁹⁹ |
| Left | Mann–Whitney U | Head-contact proportion | 165 | 168 | 27720.00 | — | 2.61566 × 10⁻⁶⁹ |
| Left | Chi-square | Dominant subfield | 165 | 168 | 329.80 | 5 | 3.89974 × 10⁻⁶⁹ |
| Right | Fisher exact | Head- vs body-dominant map | 166 | 167 | ∞ | — | 1.85967 × 10⁻⁹⁵ |
| Right | Mann–Whitney U | Head-contact proportion | 166 | 168 | 27888.00 | — | 3.59302 × 10⁻⁷¹ |
| Right | Chi-square | Dominant subfield | 166 | 168 | 322.56 | 5 | 1.41047 × 10⁻⁶⁷ |
| Pooled | Fisher exact | Head- vs body-dominant map | 331 | 335 | ∞ | — | 1.18243 × 10⁻¹⁹⁴ |
| Pooled | Mann–Whitney U | Head-contact proportion | 331 | 336 | 111216.00 | — | 1.08334 × 10⁻¹³⁸ |
| Pooled | Chi-square | Dominant subfield | 331 | 336 | 651.61 | 5 | 1.42012 × 10⁻¹³⁸ |

***Note****.* Anterior and posterior routes were compared separately by side and after pooling sides.

**Supplementary Table S7. Within-Profile Cluster-Mass-Corrected Delay-Discounting Associations**

| **Pathway** | **Hemisphere** | **n** | **Nodes** | **Mass** | **Peak** | **β** | **t** | **pFWER** |
| --- | --- | --- | --- | --- | --- | --- | --- | --- |
| Inferior VTA–NAc tract | Left | 173 | 0–47 | 113.27 | 39 | .196 | 2.65 | .0106 |
| Superior VTA–NAc tract | Right | 173 | 19–46 | 69.22 | 41 | .207 | 2.73 | .0191 |
| VTA–amygdala | Left | 168 | 8–29 | 54.84 | 24 | .218 | 2.90 | .0392 |
| VTA–posterior HPC | Left | 168 | 0–17 | 43.90 | 13 | .217 | 2.94 | .0434 |

***Note.*** Positive β indicates higher NDI with steeper discounting after sign reversal of area under the curve. Error control applies across 100 nodes within each tract-by-hemisphere profile.

**Supplementary Table S8**

*Descriptive Statistics for Participant Mean Streamline Length by Pathway and Hemisphere*

| **Pathway** | **Side** | ***n*** | ***M (mm)*** | ***SD*** | **Median (mm)** | **Range (mm)** | **CV** |
| --- | --- | --- | --- | --- | --- | --- | --- |
| Inferior VTA–NAc tract | Left | 173 | 21.91 | 1.36 | 21.78 | 18.66–25.33 | .062 |
| Inferior VTA–NAc tract | Right | 173 | 22.24 | 1.38 | 22.26 | 18.49–25.76 | .062 |
| Superior VTA–NAc tract | Left | 173 | 28.38 | 1.82 | 28.34 | 21.18–33.08 | .064 |
| Superior VTA–NAc tract | Right | 173 | 28.87 | 1.95 | 28.95 | 21.18–33.72 | .068 |
| VTA–amygdala | Left | 168 | 31.20 | 1.38 | 31.03 | 28.39–34.63 | .044 |
| VTA–amygdala | Right | 168 | 31.14 | 1.16 | 30.96 | 28.86–35.62 | .037 |
| VTA–anterior HPC | Left | 166 | 40.30 | 2.72 | 40.24 | 34.71–48.14 | .068 |
| VTA–anterior HPC | Right | 166 | 41.39 | 2.82 | 41.15 | 33.46–53.25 | .068 |
| VTA–posterior HPC | Left | 168 | 47.27 | 3.53 | 47.55 | 39.40–56.76 | .075 |
| VTA–posterior HPC | Right | 168 | 46.15 | 3.07 | 46.36 | 37.74–53.28 | .067 |
| HPC–NAc–VP | Left | 166 | 92.66 | 4.22 | 92.83 | 75.67–100.12 | .046 |
| HPC–NAc–VP | Right | 166 | 93.68 | 3.03 | 93.96 | 81.47–100.51 | .032 |
| VP–VTA | Left | 170 | 17.60 | 1.07 | 17.46 | 15.46–21.96 | .061 |
| VP–VTA | Right | 170 | 19.33 | 1.26 | 19.34 | 15.98–22.53 | .065 |

*Note.* Each participant’s value was the ordinary unweighted mean length of accepted streamlines in a valid native-space tractogram. M, SD, median, range, and CV summarize these participant-level mean lengths. CV = coefficient of variation (SD/M); HPC = hippocampus; NAc = nucleus accumbens; VP = ventral pallidum; VTA = ventral tegmental area.
